# Multiscale modeling of cell–substrate adhesion dynamics: effects of integrin activation, clustering and internalization

**DOI:** 10.1101/2025.09.05.674169

**Authors:** Huiyan Liang, Wei Fang, Xindong Chen, Bo Li, Xi-Qiao Feng

## Abstract

Cell adhesion is a fundamental biological process that governs cell proliferation, differentiation, migration, and tissue development. Cells adhere to the extracellular matrix through specialized transmembrane proteins, whose structures and functions are well studied. However, how mechanical, chemical, and biological factors interact to regulate these proteins and hence to shape cross-scale adhesion dynamics from molecular clustering to cellular migration remains unclear. Here, we propose a multiscale mechano-biochemical coupling framework to investigate the dynamics of cell–substrate adhesions, incorporating comprehensive molecular steps in the integrin life cycle, including activation, clustering, signal transduction and internalization. Our model elucidates the roles of caveolin transport and actin flow in modulating integrin dynamics and FA morphology. We identify the antagonistic interplay between integrin internalization and clustering that governs cross-scale adhesion dynamics. Furthermore, our model quantitatively demonstrates how the substrate stiffness regulates the integrin clustering size and internalization rate. These findings provide mechanistic insights into the regulation of cell migration, particularly the transition between durotaxis and negative durotaxis, driven by intracellular and extracellular microenvironmental factors. Our model offers an effective framework for understanding the cross-scale regulation process of cell adhesion involved in physiological and pathological activities, such as stem cell differentiation and cancer metastasis.

## INTRODUCTION

Cells interact with and adhere to each other or the extracellular matrix (ECM) through a variety of transmembrane molecules, such as integrins, cadherins and selectins. Among these, integrins play a central role in the formation and dynamics of focal adhesions (FAs), which enable cells to sense and respond to microenvironmental cues. The morphology of FAs depends on the formation and mature of integrin adhesion complexes (IACs). Serving as hubs of complex molecular signaling networks, IACs link the cytoskeleton to the ECM, thereby establishing force transmission and mechanoregulation pathways that respond to physical and biochemical stimuli from both intracellular and extracellular environments (1). The mechanobiochemical coupling dynamics of IACs regulate of cell adhesion behaviors, which are critical for the processes such as morphogenesis (2–5), tumor metastasis (6, 7), and epigenetic regulation (8–10). Increasing efforts have been directed towards the mechanical regulation of integrin trafficking (3, 11). These experimental results show that mechanical and biochemical cues significantly influence the vesicular trafficking of integrin, and contribute to the regulation of the cell–substrate adhesion. However, the underlying molecular mechanisms remain inadequately understood.

Integrin serves as the key regulatory point of vesicular trafficking mediated by adaptor proteins, including clathrin (12, 13), caveolin-1 (CAV-1) (14), and clathrin- independent carriers like glycosphingolipid (14, 15). It can traffic via membrane uptake and outward delivery, i.e., endocytosis and exocytosis. Under mechanical stimuli such as shear, stretch or osmotic shock, cell membrane tension increases, suppressing the flat-to-curved transition during endocytosis mediated by clathrin (16), caveolin (17) and other carrier components such as CLIC/GEEC (14). The mechanosensitivity and responsiveness of these carriers are critical for vesicular trafficking of integrins. Recent studies highlight that mechanosensitive integrin trafficking significantly influences multiscale adhesion dynamics and various cellular responses. At the adhesion cluster scale, CAV-1 contributes significantly to FA morphology and turnover dynamics (18– 20). The mechanobiochemical properties of extracellular matrix and intracellular cytoskeleton are key regulatory factors of multiscale adhesion dynamics. While it was hypothesized that soft substrates inhibit integrin activation through membrane fluctuations (21, 22), Du *et al*. experimentally demonstrated a contrasting effect of substrate stiffness on integrin activation (23). They proposed an alternative mechanism, where CAV-1-mediated β1 integrin internalization overcompensates, leading to suppressed adhesion and enhanced neurogenic lineage specification on soft substrates (23). The cytoskeleton is a highly complex and interconnected system closely linked to cell migration in processes such as cancer metastasis, immune surveillance, and embryonic development (1, 24, 25). Besides, the cytoskeleton, particularly actin and microtubules, actively interacts with the endocytosis and exocytosis process to regulate integrin dynamics as well as nanoparticle transport (13, 17, 26). Insights into endocytosis mechanisms have been instrumental in designing efficient nanoparticle carriers for mRNA vaccines, drug delivery, and gene therapy, aided by spatiotemporal simulations using coarse-grained molecular dynamics and theoretical models (27–30). Despite these advancements, the specific mechanisms by which mechanical stimuli influence integrin trafficking and regulate cell adhesion dynamics remain poorly understood.

Theoretical and numerical research on the regulation of integrin trafficking on cell adhesion have largely focused on the molecular scale. The mechanical principles of FAs in cell–matrix interface has been systematically investigated using a coupled stochastic–elastic framework by Gao *et al*. (31). Their results reveal factors that promote the stability of FAs, including stiff substrate, intermediate adhesion size, stiffening and low-angle pulling of cytoskeleton. Based on this model, Qian and Gao further found that soft matrices suppress the cooperative effect among open receptor– ligand bonds (32). Gong *et al*. examined the motor clutch dynamics between cells and viscoelastic substrates using analytical methods and Monte Carlo simulations (33). Theoretical analysis by Du *et al*. shows that integrin–ligand binding is more prone to rupture on a soft substrate, providing insights into enhanced integrin internalization (23). Xu *et* al. proposed a biomechanical model to study how the mechanical regulation of integrin activation and internalization by ECM elasticity (34). In our recent work (35), we developed a mechano-biological coupling model to study the multiscale adhesion process from integrin clustering to FA formation. While our model elucidates the regulation mechanisms of FA morphology, motion and assembly, and cell–substrate interface force transmission, some key mechanisms remain unexplained. For example, the model does not consider the impact of integrin trafficking on cell adhesion. To better understand integrin-trafficking-related adhesion dynamics and cellular responses to intra- and extra- cellular microenvironmental factors, models capable of capturing the molecule–cluster–interface–scale–spanning dynamics regulated by mechanochemical coupling mechanisms are urgently needed but remain lacking.

In this study, we establish a multiscale dynamic model of cell–substrate adhesion that incorporates key processes of the integrin life cycle, including activation, clustering and internalization. This paper is outlined as follows. First, we formulate the bond dynamic model of integrin internalization, and incorporate it to the numerical framework of our previous multiscale model of the cell–substrate interface. Using this model, we theoretically explore the multiscale regulation of integrin internalization on the cell–substrate adhesion dynamics in response to the substrate rigidity, and validate the model based on the experimental results. Next, we perform numerical analyses to examine how the critical steps in integrin life cycle, i.e., integrin internalization and integrin clustering, regulate the adhesions at multiple scales. Finally, we conduct numerical simulations to elucidate mechanobiochemical regulation on FA morphology and cell migration, influenced by both the extracellular substrate and the intracellular actin cytoskeleton.

## MODEL

Our model aims to investigate the integrin-mediated cell–substrate adhesion to uncover the multiscale mechanobiochemical coupling regulation mechanism governing cell adhesion. As illustrated in Fig. 1, our theoretical framework captures cross-scale adhesion dynamics across the structural levels of integrin molecules, FAs, and the cell– substrate interface. It also bridges these molecular and structural interactions to cellular behaviors driven by interface sensing. Integrin molecules serve as the primary receptors mediating the adhesion dynamics at the cell–substrate interface. In this model, we incorporate key steps in the integrin life cycle [Fig. 1 (b)], which typically progresses through activation, ligand binding, clustering, and signal transduction, in a sequential order. Simultaneously, integrin internalization through endocytosis occurs and, meanwhile, internalized integrins are recycled back to membrane. Based on the previous work (35), we extend the model to incorporate integrin trafficking, thereby developing a comprehensive multiscale mechanobiochemical framework for decoding cell–substrate adhesion.

**Fig. 1.**
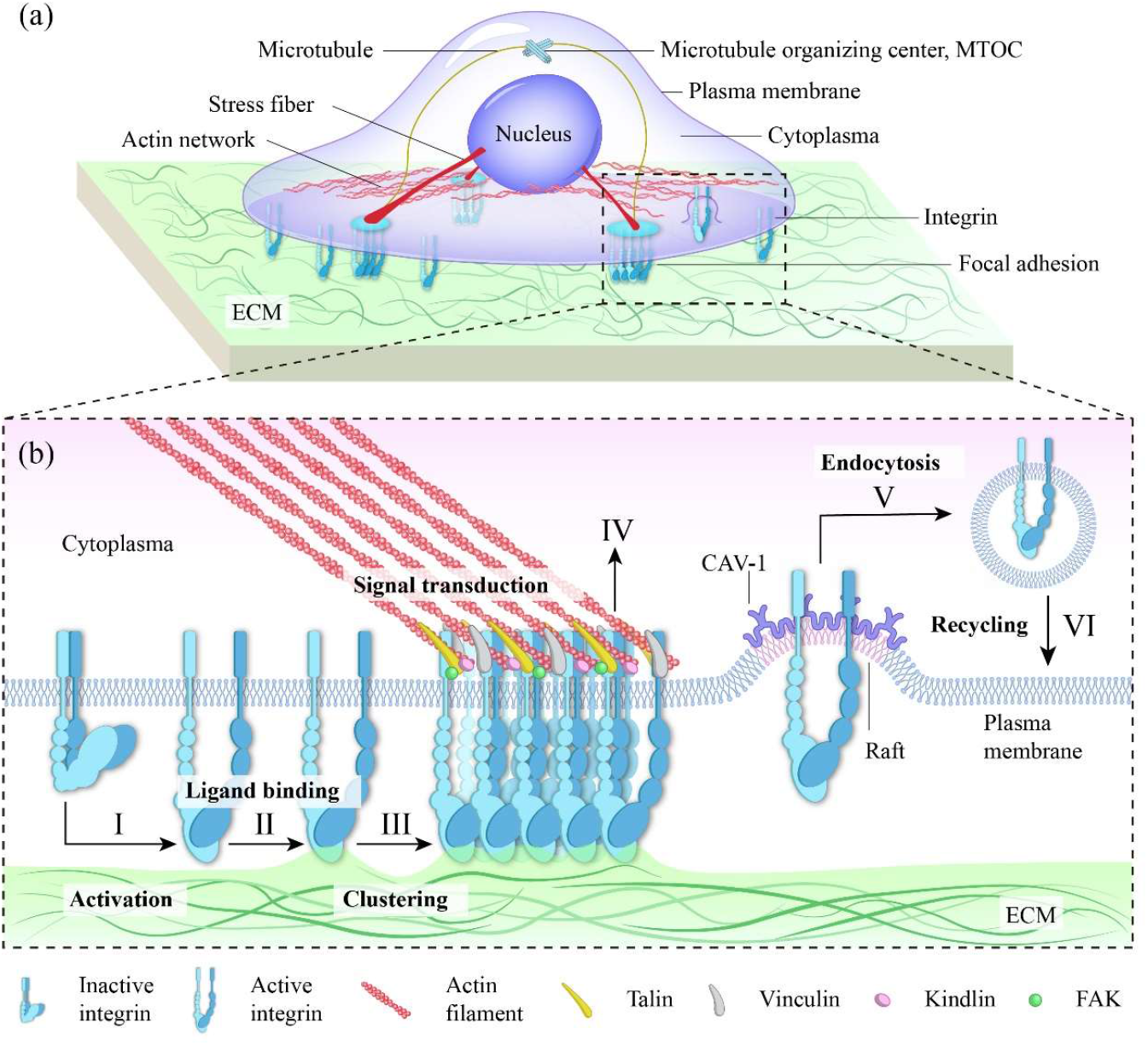
Schematics of cross-scale dynamics in cell–substrate adhesions. (a) A single cell interacting with the ECM through the cell–substrate interface. (b) Key steps of the integrin life cycle at the cell–substrate interface.

### Bond dynamics of integrin internalization

Vesicular trafficking of integrins depends on CAV-1-mediated internalization. CAV-1 consists of an N-terminal tail and a C-terminal segment connected by a sequence that includes a scaffolding domain and an intramembrane domain (36–39). With this hairpin structure, CAV-1 is anchored stably to the inner surface of the cell membrane (38, 39). Caveolins aggregate into multimolecular complexes on the membrane, driven by interactions involving the N-terminal domain (40, 41), which may also include specific C-terminal-to-C-terminal attractions (42). CAV-1 drives membrane invaginations to form caveolae, facilitating integrin internalization and promoting cellular mechanoadaptation (43, 44).

We develop a bond dynamics model for integrin internalization. It is assumed that liganded or clustered integrins are allowed to be internalized. When the internalized integrin is trafficked into the cytoplasm, the corresponding integrin–ligand (IL) and integrin–integrin (II) bonds break, and reversible association is not allowed. Since no external forces act on the internalized integrins to extend their conformations, we consider them to be in an inactive state. There are 

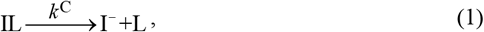

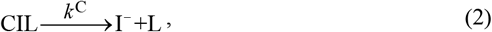

where *k*^C^ is the integrin internalization rate. Denote I^−^ as the inactive integrin, and L as the ligand. IL and CIL represent ligand-bound integrins and clustered integrins, respectively. To simplify the modeling of adhesion–actin flow engagement, clustered integrins (CIL) are assumed to bind instantaneously with actin filaments upon clustering. Therefore, CIL is considered equivalent to ACIL (actin-bound clustered integrins) throughout this work. Activated integrins that are not bound to ligands can also undergo internalization within a cholesterol-enriched membrane microdomain, known as lipid raft (45, 46). We assume that the internalization rate of these integrins is equal to that of the liganded integrins, given by

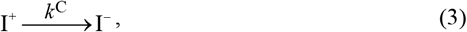

where I^+^ represents the activated integrin.

Based on previous studies (34, 39), the association between integrins and CAV-1 induces membrane invagination, generating a force that acts on the integrins. As shown in Fig. 2(a), *F*_C_ represents the characteristic force driving integrin internalization, which facilitates the detachment of integrins from the substrate. This characteristic force is not directly included in the lateral motion equation of integrins, but rather enters the model as a regulator of the integrin internalization rate, reflecting its role in facilitating integrin unbinding during CAV-1-mediated endocytosis. In this sense, the mechanical effect is captured through a force-dependent reaction kinetics term, rather than as an explicit mechanical force in the motion dynamics. The greater the force exerted by CAV-1, the more readily integrin internalization occurs. The internalization rate *k*^C^ can thus be expressed as

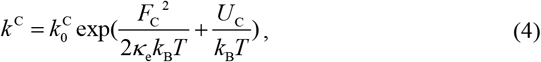

where *U*_C_ is a negative parameter representing the energy released via IL binding, 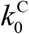 denotes the base internalization rate of integrin, and *κ*_e_ represents the effective stiffness of the integrin–substrate (IS) link. *k*_B_ and *T* denote Boltzmann’s constant and temperature, respectively. Based on the series connection between the IL bonds and the substrate surface which together form the IS chain, *κ*_e_ satisfies the following relation

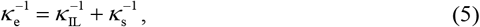

where *κ*_IL_ and *κ*_s_ represent the stiffness of the IL bonds and the effective surface stiffness of the substrate, respectively. *κ*_s_ can be expressed as (23)

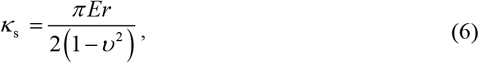

where *E* and *υ* represent the Young’s modulus and Poisson’s ratio of the substrate, respectively. *r* denotes the radius of integrins. Moreover, integrin trafficking via endocytosis is a broadly conserved feature across integrin subtypes, although the specific routes vary depending on the subtype—for instance, α5β1 undergoes clathrin-mediated endocytosis. While the current bond dynamics model is developed in a CAV- 1-related context, it can be readily adapted to different integrin subtypes.

**Fig. 2.**
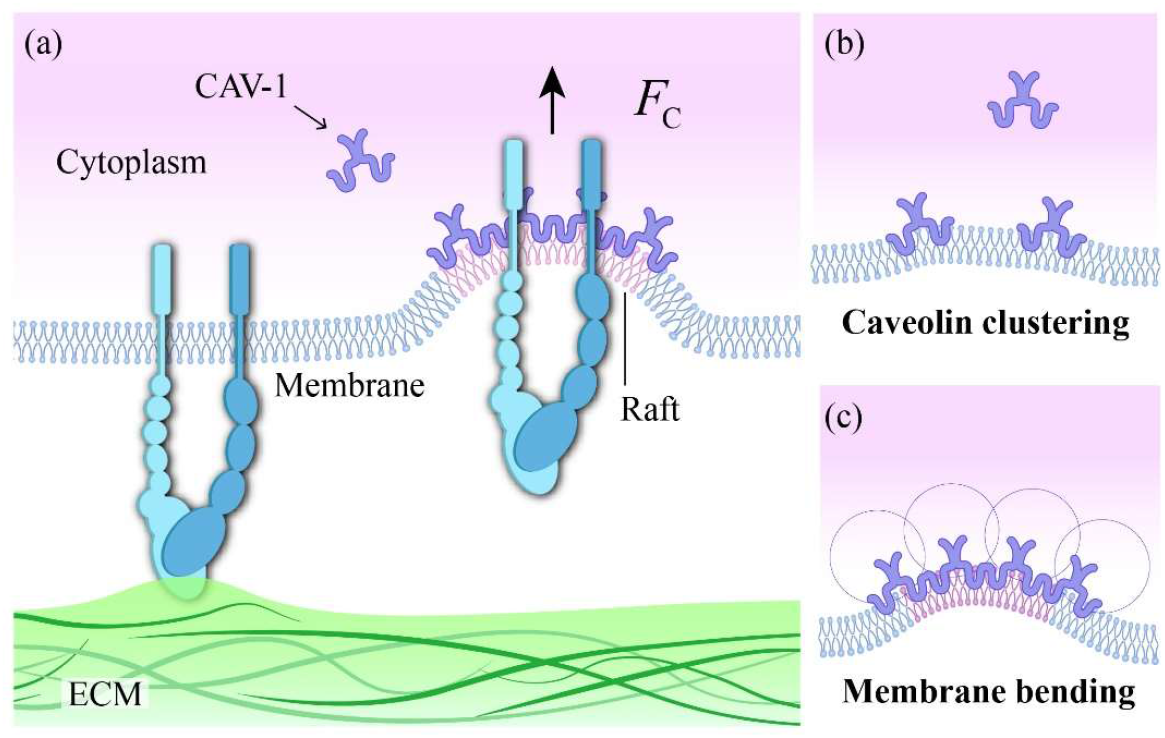
Schematics of (a) caveolin-mediated integrin internalization, (b) caveolin clustering driven by residues of the N-terminal located close to the membrane, and caveolin inducing plasma membrane invaginations.

### Multiscale adhesion model of integrin internalization

We incorporate the bond dynamics model of integrin internalization into the multiscale numerical framework of adhesion dynamics proposed in our earlier work (35) [Fig. 3(a)].

**Fig. 3.**
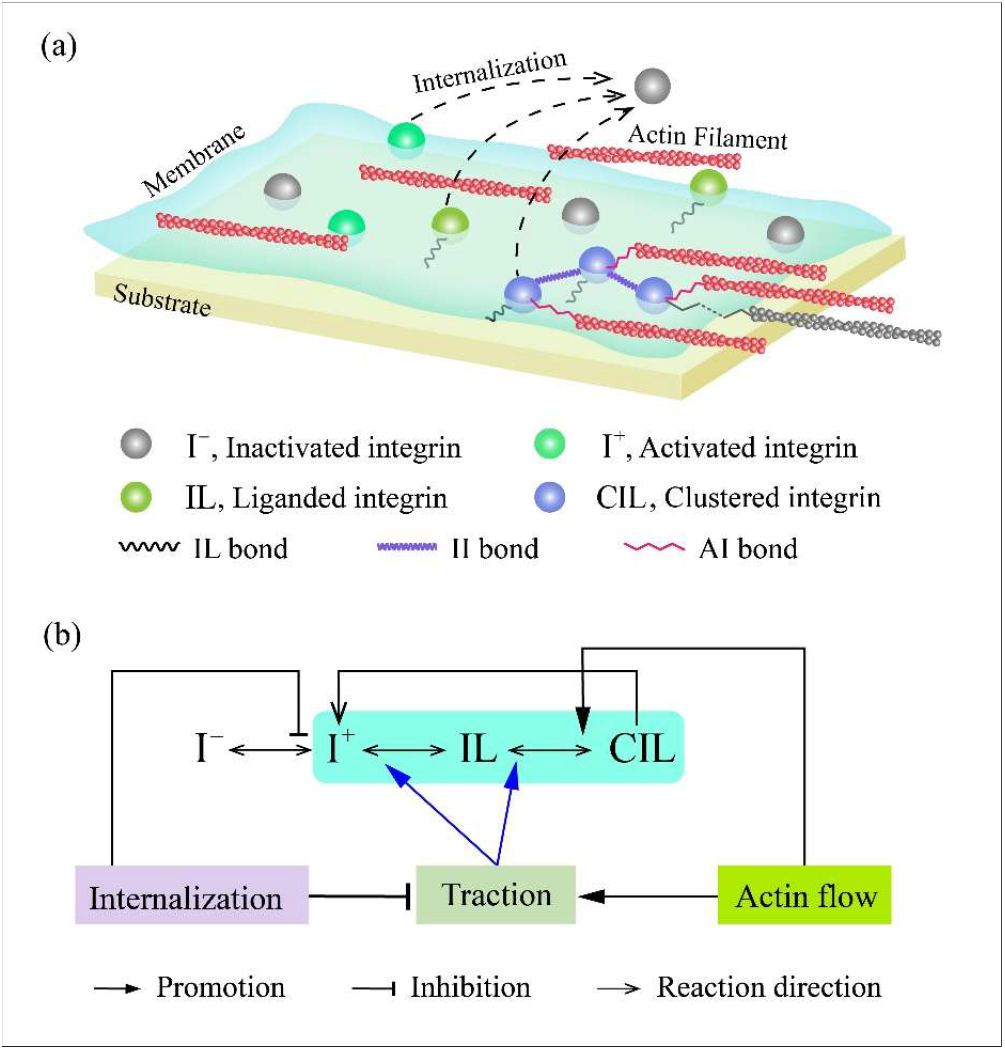
(a) Schematic of the life cycles of integrins including activation, clustering and internalization. CIL with AI bond is actin-bound clustered integrin (ACIL). (b) Schematic of the mechanobiochemical coupling regulations of the multiscale cell– substrate adhesion model.

Integrins are modeled as mutually repulsive diffusing transmembrane particles subjected to intra- and extra- cellular interactions [Fig. 3(a)]. Due to their embedding in a viscous phospholipid bilayer, the motion of integrins is regarded to be overdamped and satisfies (47), 

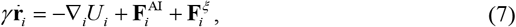

where **r**_*i*_ (*t*) and 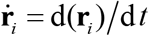 denote the real-time position of integrin *i* and the corresponding time derivative, respectively, both as vectors in three-dimensional Euclidean space. *γ* is the damping coefficient of integrins. The forces exerted on integrin *i* include the cytoskeletal force 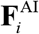, the stochastic force 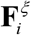, and the configurational force −∇_*i*_*U*_*i*_. The configurational potential *U*_*i*_ of integrins *i* 

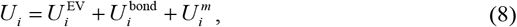

where the contributions of volume exclusion interaction 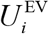, intermolecularinteraction via bond association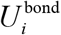, and membrane friction 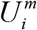, are considered.

Here, we consider constant-rate activation and deactivation of integrins 

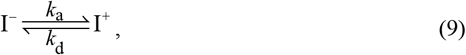

where I^−^ and I^+^ denotes the inactivated and the activated integrins, respectively. *k*_a_ and *k*_d_ represent the reaction rate of activation and deactivation. The ratio of them is determined by the activation energy *E*_a_, 

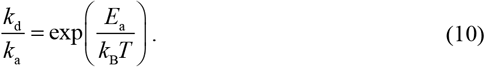

The assembly of cell–substrate adhesions governed by both in-plane and out-of-plane bond dynamics. The in-plane dynamics refers to integrin clustering [Eqs. (12) and (13)], while the out-of-plane dynamics encompasses ligand binding [Eq. (11)], interactions with actin cytoskeleton [Eqs. (14) and (15)], and integrin internalization [Eqs. (1–3)]. These are given as 

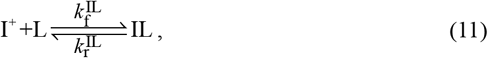

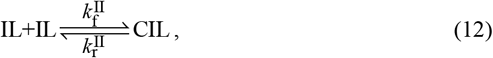

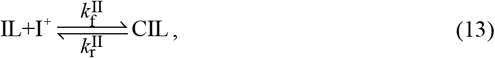

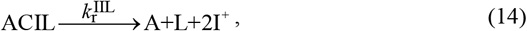

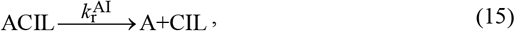

where the forward and reverse reaction rates of the corresponding reactions are indicated by the above and below reaction arrows, respectively; and the reactants and products are indicated in Fig. 3.

In the reverse reactions of Eqs. (11–13), the bond rupture rates of IL and II bonds are governed by the Bell model (48) 

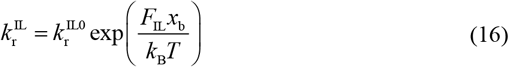

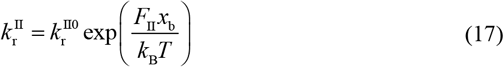

where 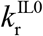 and 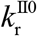 are the base rupture rates of IL and II bonds. *F*_IL_ and *F*_II_ denote the IL and II bond forces, respectively. *x*_b_ is the transition distances of IL and II bonds, and is positively correlated to the mechanosensitivity of IAC bond dynamics.

Under the tension, mechanosensitive adaptor proteins of IACs, e.g., talin and vinculin could reinforce the actin–adhesion association, initiating adhesion-localized actin bundling which facilitates integrin clustering (49–51). To capture this mechanosensitive process, we consider the II clustering rate 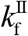 in Eqs. (12–13) upregulated by actin–integrin (AI) bond force *F*_AI_ 

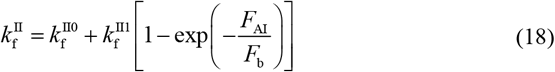

where 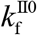 and 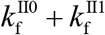 denote the non-load and the maximum binding rate of II bonds, and *F*_*b*_ is the characteristic force describing the mechanosensitivity of 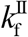 to *F*_AI_.

## NUMERICAL SCHEME

The multiscale model is developed using a coarse-grained particle-based framework, and is solved numerically due to challenges of obtaining an analytical solution. A three-dimensional computational representative volume element (RVE) domain is constructed with dimensions of *L*_*a*_ × *L*_*a*_ × *h*_m_, where the in-plane dimension *L*_*a*_ is set to 1.5 μm. Periodic boundary conditions in the *x*–*y* plane is incorporated to the RVE. Initially, ligand molecules are uniformly distributed on the substrate surface (plane *z* = 0) within the RVE domain, while integrins are confined to the membrane plane (*z* = *h*_m_). The output, including coordinates and states of all integrins, bond connections, and tension vectors, are recorded every 1000 steps. At each simulation step, the integrin states are updated using the Gillespie algorithm based on stochastic chemical reactions, then the motion equation of integrins [Eq.(7)] is solved using the finite difference method. Time steps were selected based on convergence analysis and adjusted according to case-specific physical parameters, and a simple adaptive time-stepping control was implemented as a safeguard to enhance the robustness of the numerical scheme.

In simulations, the real-time proportions of integrins with IL bonds and II bonds *P*_IL_ (*t*) and *P*_II_ (*t*) are tracked, where *t* represents time. To quantitatively capture the spatial–temporal evolution of the dynamics and mechanics of adhesion clusters at the cell–substrate interface, formed adhesion clusters are identified using the density-based spatial clustering of applications with noise (52). Then, the number of formed adhesion cluster at time *t, n*_clu_ (*t*), is tracked, and the cluster density *ρ*_clu_ (*t*) can be obtained as 

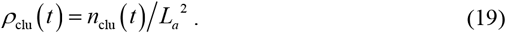

The cluster averaged values of cluster diameter Φ_clu_ (*t*), cluster integrin number 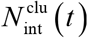, and cluster eccentricity *e*_*x*to*y*_ (*t*) at time *t* are calculated respectively as follows: 

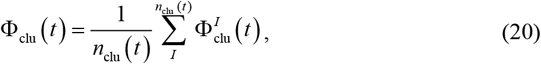

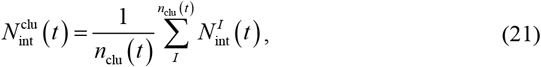

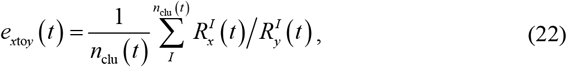

where 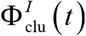 and 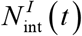 are the diameter and assembled integrin number of the cluster *I*, respectively. In Eq.(22), 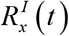 and 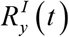 denote the mean spacings in the *x*- and *y*-directions, respectively, between each cluster centroid and its farthest integrin molecule, averaged over all adhesion clusters in the computational domain at a given time.

To characterize force transmission at the at the cell–substrate interface, the **σ**^if^ (*t*), components of interface stress **σ**^if^ (*t*) in the out-of-plane [i.e., 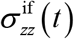 in the *z* direction] and in-plane directions [i.e., 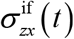 and 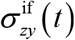 in the *x* and *y* directions, respectively] are calculated in every 20000 steps. For instance, the interface stress component aligned with the actin flow, 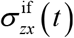, is defined by 

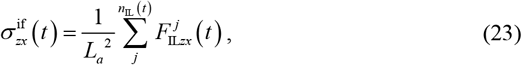

where *j* is the index of the IL bond, *n*_IL_ (*t*) is the number of IL bonds, and 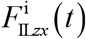 represents the *x*-component of bond force of the IL bond *j*. By an overbar, 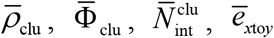, and 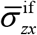 indicate the equilibrium values of are obtained by averaging over at least one time period at equilibrium state, respectively. For example, 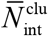 represents the average cluster integrin number at equilibrium state. These statistical quantities re summarized in Table 1.

**Table 1.** Nomenclature

| Averaged parameters at equilibrium state | Notation | Units |
| --- | --- | --- |
| Proportion of integrins with IL bonds | $\bar{P}_{\text{IL}}$ | 1 |
| Proportion of integrins with II bonds | $\bar{P}_{\text{II}}$ | 1 |
| Cluster density | $\bar{\rho}_{\text{clu}}$ | $\mu\text{m}^{-2}$ |
| Cluster integrin number | $\bar{N}_{\text{int}}^{\text{clu}}$ | 1 |
| Diameter of clusters | $\bar{\Phi}_{\text{clu}}$ | nm |
| Eccentricity of clusters | $\bar{e}_{\text{xtoy}}$ | 1 |
| Interface stress tensor | $\bar{\sigma}^{\text{if}}$ | Pa |

## MODEL VALIDATION

The mechanical regulation of integrin internalization plays an important role in FA dynamics. To validate the multiscale model that incorporates integrin internalization, we simulate the substrate stiffness-sensing behavior of cell–substrate adhesion and compare the results with relevant experimental data.

To investigate adhesion dynamics and force transmission behavior at the cell– substrate interface, we employ the multiscale model proposed in Section 2.2. Numerical simulations are conducted using an interfacial representative volume element model with in-plane dimensions of 1.5 μm, where actin flow is initiated at a velocity of 10.0 nm/s from the outset. The adhesion evolution process is examined under a constant II binding rate. As shown in Fig. 4(a), under the influence of the integrin internalization, the average number of molecules per adhesion cluster 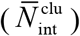 and the average cluster density at the equilibrium state 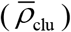 increase significantly with increasing Young’s modulus *E* of the substrate in the low-to-medium modulus range (10–50 kPa) and tend to plateau in the high modulus range (100–1000 kPa). These overall trends of FA dynamics in response to the substrate stiffness resemble S-shape curves. The predicted FA scale 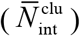 as a function of substrate stiffness shows good qualitative agreement

**Fig. 4.**
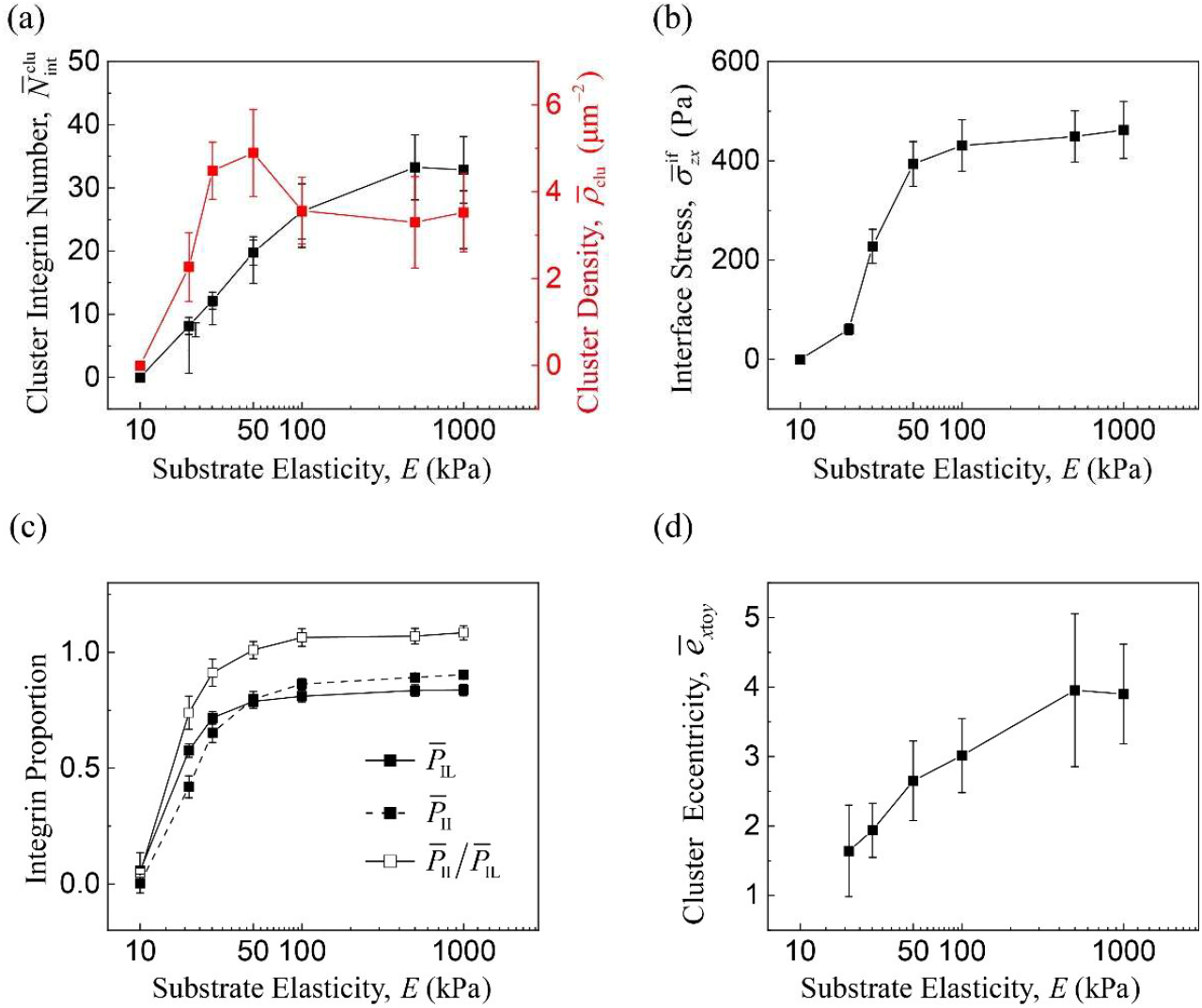
Numerical results from multiscale dynamic model considering the integrin internalization: the mean values of (a) the number of molecules within adhesion clusters and cluster density, (b) interface stress, (c) proportion of ligand-bound integrins, clustered integrins, and the ratio of clustered integrins to ligand-bound integrins, and (d) cluster eccentricity at equilibrium state. In simulations, we take 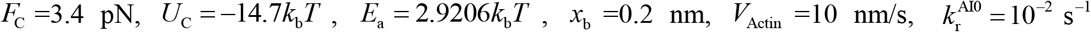, and *E_II_* =3.9889 *k_b_T*, considering constant-rate II bonding processes.

with the experimentally observed trend of increasing FA sizes (20, 53) (Fig. 5). Moreover, the predicted threshold of the Young’s modulus *E* required to form significant adhesion clusters is of the same order of magnitude as the experiment data (53). Note that the absolute cluster sizes predicted by the model differ from experimentally measured focal adhesion sizes by approximately an order of magnitude. This discrepancy arises from the coarse-grained nature of the model, where each simulated particle represents a group of molecules, and from differences in how cluster size is defined in simulation versus experiment. Nevertheless, these results demonstrate that our model the model successfully captures the mechanical regulation of integrin internalization on the FA dynamics, validating its effectiveness. Furthermore, the eccentricity of FA increases monotonically with the substrate stiffness, showing an upward trend gradually moderating and reaching a plateau in the range of 100–3000 kPa [Fig. 4(d)]. This result suggests that, with adhesion assembly being promoted by the mechanical regulation of integrin trafficking, the anisotropic growth of adhesion clusters is correspondingly facilitated through actin flow–mediated physical transport. On a substrate with *E*=10 kPa, almost no adhesion clusters are visible, while small, sparsely distributed adhesion clusters appear at *E*=20 kPa. As *E* increases, the scale of adhesion clusters augments, and the polarity along the actin flow direction become internalization rates more pronounced, consistent with the orientation of highly polarized FAs observed experimentally [Fig. 5(b2)].

**Fig. 5.**
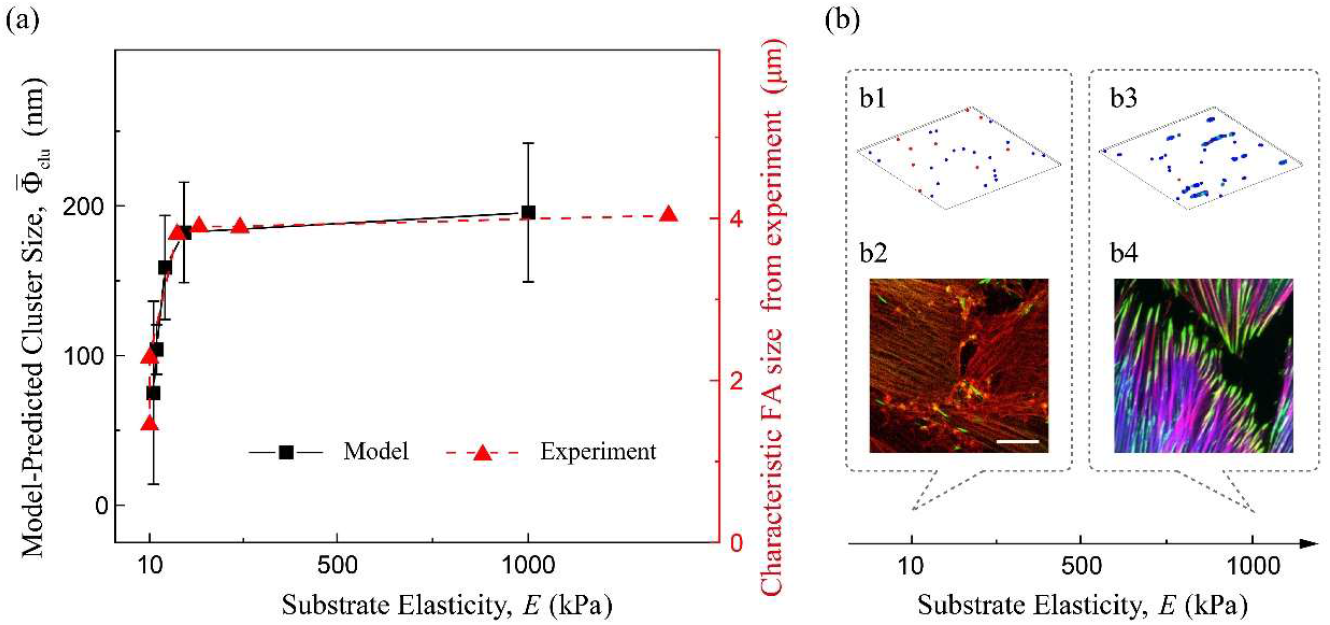
Comparison of numerical results for FA size regulated by Young’s modulus of the substrate with experimental measurements (53). Panels b1–4 show representative adhesions on substrates with low and high substrate stiffness, respectively. Among them, b1 (*E*=10 kPa) and b3 (E=1000 kPa) present simulated integrin clustering, whereas b2 and b4 display fluorescence microscopy images of actin filaments and focal adhesions adapted from (53). In the experiment of Goffin *et al*. (53), REF-52 myofibroblasts were fixed and stained for vinculin, α-SMA, and F-actin, which were visualized in green, blue, and red, respectively (a2 and b2), after 12 hours of culture on PDMS substrates with Young’s moduli of 9.6 kPa (a2) and 780 kPa (b2). In simulations, we take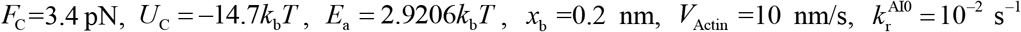, and *E*_II_ =3.9889 *k*_*b*_ *T*, considering constant-rate II bonding processes.

Consistent with the observed trends for 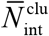 and 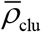, the equilibrium-state proportions of ligand-bound and clustered integrin molecules (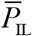 and 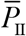) at the molecular scale, as well as the interfacial traction stress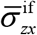, also exhibit S-shape trends with an increasing Young’s modulus *E* [Fig. 4(c)]. These trends reflect an initial rapid rise followed by a plateau. Among these, the change in 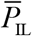 is more pronounced than that in 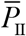 as substrate stiffness increases, suggesting that the mechanical regulation of integrin internalization exerts a stronger inhibitory effect on adhesion clustering than on adhesion formation. The interfacial traction stress is primarily governed by the out-of-plane adhesion dynamics (i.e., IL bonding and AI bonding) at the cell–substrate interface. Then, increasing *E* upregulate the interfacial traction stress.

The sharp rise of 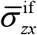 when *E* increases from 10 kPa to 50 kPa is attributed to attenuated integrin internalization, which facilitates the formation of stable adhesion clusters for effective force transmission. The subsequent plateauing of 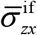 and 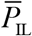 after E reaching ∼50 kPa [Fig. 4 (b)] reveals a balance of reduced integrin internalization and increased IL bond rupture, leading to saturated force transmission.

## RESULTS AND DISCUSSION

We have established the theoretical model and numerical methods to depict the dynamic process of integrin internalization. Subsequently, we examine the mechanical regulation of integrin internalization on the cell–substrate adhesion dynamics across multiple structural scales and validate our model by comparing numerical results with experimental data.

As highlighted in the introduction, current understanding and its cross-scale impact on cell adhesion remain limited. Therefore, in this section, we employ the multiscale dynamic model of integrin internalization to explore the mechanosensing and coordinating mechanisms that govern adhesion behaviors. Herein, we focus on how the mechanosensitivity of key molecular steps in the integrin life cycle, in conjunction with intra- and extracellular factors, regulates these processes.

### The effect of integrin internalization on adhesion dynamics

Within the IACs molecular network, certain components play distinct roles in regulating adhesion dynamics. For instance, CAV-1 mediates the mechanical regulation on adhesion dynamics by controlling the out-plane membrane uptake of integrins. Conversely, components such as talin regulate in-plane dynamics which determines the lateral assembly of adhesion clusters. In this subsection, we quantitatively analyze the effect of clustering rate on the multiscale adhesion dynamics under the process of integrin internalization.

In the simulation, we consider the multiscale spatiotemporal evolution of the cell– substrate adhesion on a substrate with *E* = 100 kPa under the actin shear flow with a velocity of 10 nm/s. To investigate the effect of integrin II bond formation rates (i.e., integrin clustering rate), a series of dimensionless II bond binding energy 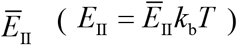 are used for numerical simulations. Through these simulations, a quantitative relationship between multiscale adhesion dynamics and integrin clustering rate is established (Fig. 6).

**Fig. 6.**
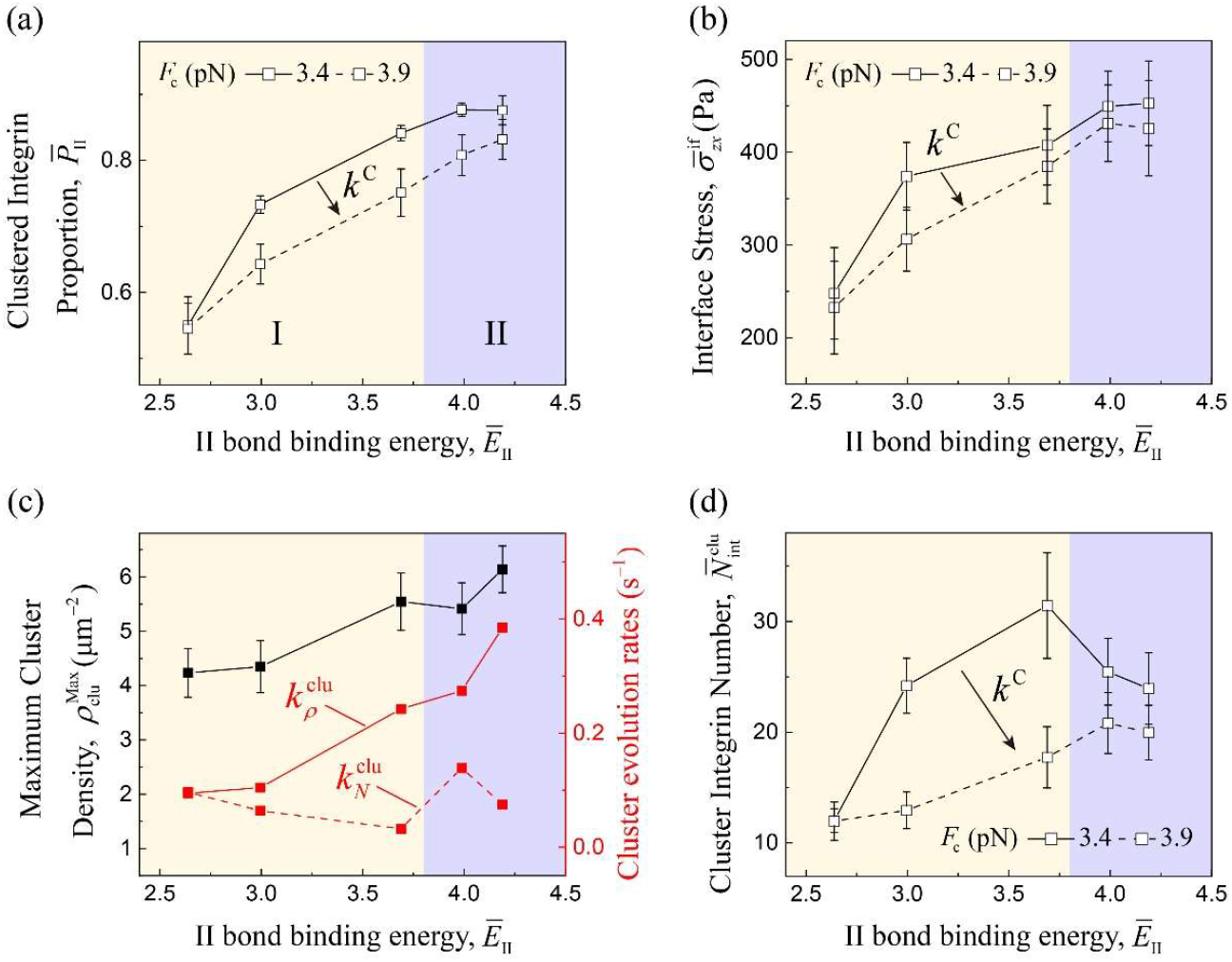
Effect of II bond binding energy on the multiscale dynamics of cell–substrate adhesions: the mean values of (a) the proportion of ligand-bound integrins and clustered integrins, (b) interface stress, (c) maximum cluster density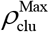, cluster nucleation rate 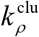 and cluster assembly rate 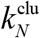, as well as (d) number of molecules within adhesion clusters at equilibrium state. The orange region and the purple region represent the stage I and II, respectively. In simulations, we take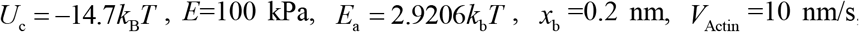, and 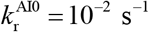, considering constant rate II bonding processes. Arrows indicate the direction of increasing integrin internalization rate.

The coupling of integrin internalization with integrin clustering rate to regulate multiscale adhesion dynamics is examined by simulating two cases: one with a low internalization rate (at *F*_c_ = 3.4 pN), and the other with a high internalization rate (at *F*_c_ = 3.9 pN). In the low-to-medium rate range (i.e., Stage I, *Ē*_II_= 2.6391−3.6889), the clustering rate promotes the adhesion dynamics of integrin molecules, cluster assembly and interface transmission. However, when reaching the high-rate range (i.e., Stage II, *Ē*_II_= 3.9889−4.1897), these monotonic positive relations begin to depart. At the molecular scale, the increasing clustering rate promotes integrin clustering (i.e., 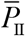) in Stage I [Fig. 6(a)]. In Stage II, however, the promotion diminishes, with 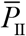 plateauing at of 87% after a minor further promotion [Fig. 6(a)]. At the interface scale, force transmission is significantly enhanced in Stage I and reaches a plateau in Stage II, corresponding to the variation in 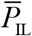 [Fig. 6(b)]. At the FA level [Fig. 6(c, d)], 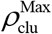 and 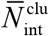 are positively correlated with 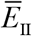 in Stage I, but in Stage II, they diverge— 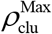 continues to increase, while 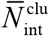 decreases. These results reveal that integrin clustering can simultaneously promote the assembly and nucleation of adhesion clusters, then leads to the nucleation– assembly competition when the molecular clustering activity goes too high. A similar competition also occurs between the rates of cluster nucleation and cluster assembly, i.e., 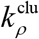 and 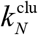 as shown in Fig. 6(c). During the full stage, in the low-internalization-rate case (i.e., *F*_c_ = 3.4 pN in Fig. 6), adhesion dynamics at the molecular, FA and interface scales in response to increasing Ē _II_initially exhibit a monotonic positive trend, followed by a saturation stage at higher rates. The downregulated cluster assembly by *Ē*_II_ in Stage II, should reflect the constraining effect of a limited pool of surface integrins due to internalization. Interestingly, however, the resultant reorganization of adhesion clusters in Stage II demonstrates a capacity to rebalance through smaller-scale and denser distributions under a nearly constant load level, as integrin clustering activity increases.

Through further comparative analysis, our simulations reveal the regulatory role of integrin internalization in multiscale adhesion dynamics. As shown in Fig. 6(a, b) and Fig. S3(b), an increase in *k*^C^ leads to reduced 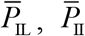 and 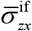, showing that integrin internalization can inhibit adhesion formation and integrin clustering at the molecular scale, as well as interface force transmission. Moreover, 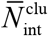 is evidently lower in the case with higher *k*^C^, exhibiting that integrin internalization can impose effective inhibition of FA assembly [Fig. 6(d)]. These results agree well with related experiments, where expression of CAV1, which is suggested to be closely related to increased overall internalization rate of integrin (23), leads to reduced integrins on the cell surface and promoted FA disassembly (23, 54). Here, we adopt the dimensionless parameter 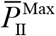, i.e., the maximal proportion of clustered integrin, as the saturation coefficient that characterizes the upper limit of integrin molecules involved in clustering. It is worth noting that the molecular-scale saturation in response to increasing *Ē*_II_ are delayed by promoted integrin internalization. These results can be attributed to the “constraining” effect of integrin internalization dynamics, which couples with the recycling dynamics through ligand binding and clustering to establish the dynamic equilibriums at the adhesion interface, thus limiting the total number of available integrin molecules. The competition between growth and nucleation of FAs can be interpreted from the cross-scale interactions between the molecular and FA levels—the FA evolution is constrained by the balance between integrin trafficking and recycling.

The effects of two key molecular-scale factors — integrin internalization and integrin clustering—on multiscale adhesion dynamics are summarized. As shown by Fig. 7, the internalization and clustering of integrins exhibit antagonistic effects in regulating the processes of endocytosis, II association and IL binding. The influence of the antagonistic mechanism is transmitted to higher structural levels, resulting in cross-scale regulation on cell–substrate adhesion behaviors. These processes ultimately determine the evolution of the focal adhesion size and distribution at the cell’s leading edge and tail.

**Fig. 7.**
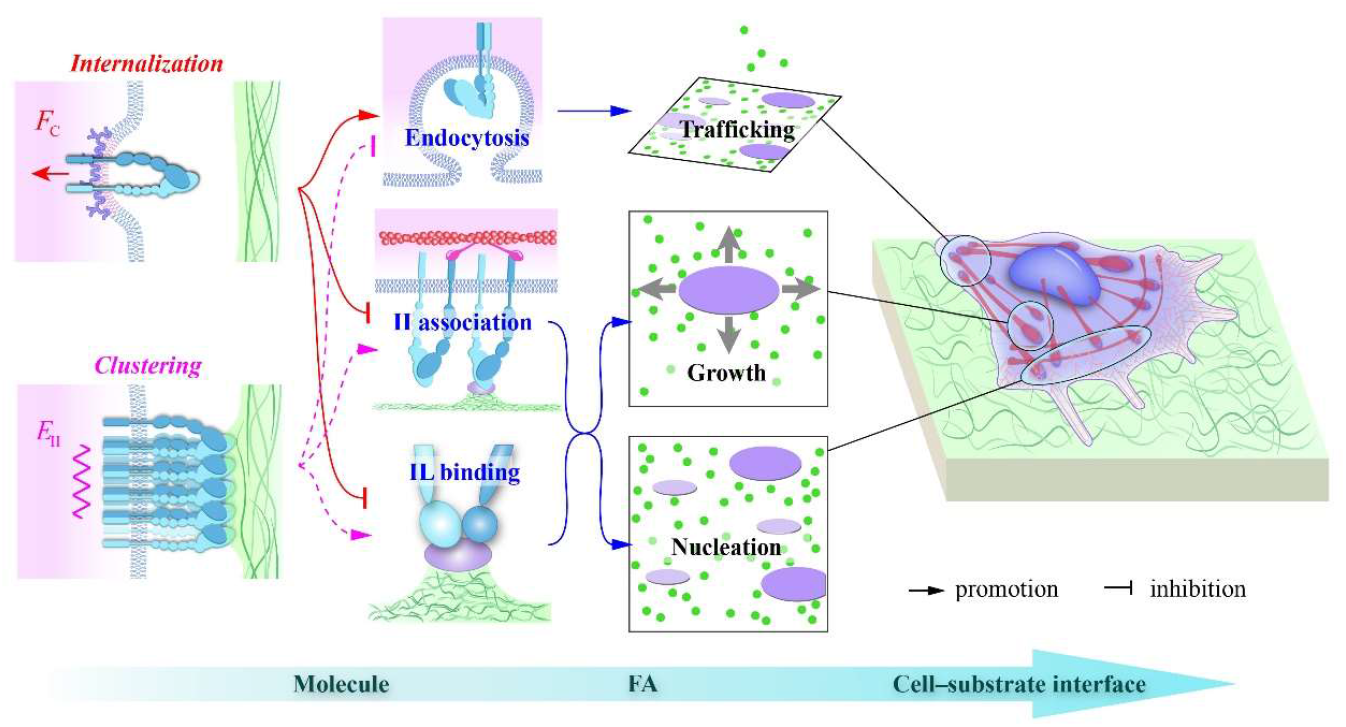
Schematic of the cross-scale adhesion regulation of *E*_II_ at the cell–substrate interface considering the integrin internalization.

### Substrate rigidity sensing of multiscale adhesion dynamics

It has been recognized that the mechanosensitivity of cell dynamics originates from the molecular transaction network that integrates mechanical information with biochemical signals (11). A key step in this process—integrin internalization, mediated by mechanosensitive proteins such as CAV-1—regulates multiscale adhesion dynamics by promoting the uptake of integrins into the plasma membrane. This subsection will examine the effect of integrin internalization on multiscale adhesion dynamics at cell– substrate interface.

We first simulated the two sets of internalization parameters shown in Fig. 8, considering constant II binding rate and force sensitivity of IAC bond rupture at anappropriate level to ensure dependable force transmission (*x*_b_ = 0.2 nm). The internalization rate–Young’s modulus curves corresponding to each group of parameters are shown in Fig. 8(a). The internalization rate increases when *F*_C_ rises from 3.4 pN (black line) to 3.9 pN (blue line) [Fig. 8(a)]. As shown in Fig. 8(b–f), the regulatory effects of the substrate stiffness on adhesion dynamics at the molecular, adhesion cluster, and interface scales are generally S-shape in both cases. On softer substrates (E= ∼20– 50 kPa), smaller and denser adhesions tend to form, while larger adhesions tend to form on stiffer substrates. The increase in internalization rate significantly reduces the assembly of adhesion clusters [Fig. 8(b)], and 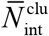 decreases by around one-third when *E* = 1000 kPa. Meanwhile, 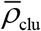 also declines when *k*^C^ gets higher when *E*≤50 kPa [Fig. 8(b)], indicating that the enhanced integrin internalization can simultaneously suppress adhesion assembly and nucleation. However, on stiffer substrates (*E*>100 kPa) [Fig. 8(b)], the enhanced integrin internalization leads to a sudden and moderate decrease of 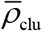 due to the limited total number of integrin molecules, which ultimately suppresses nucleation. The competition between adhesion assembly and nucleation is discussed in detail in Subsection 4.1. 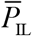 and 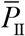 can respectively characterize the adhesion formation and the integrin clustering at the IAC molecular scale. As shown in Fig. 8(d–f), the integrin internalization process weakens on the adhesion formation, the integrin clustering, and the interfacial force transmission over the entire *E* range, among which the weakening effect is more significant on soft and medium-stiffness substrates. As the substrate stiffness increases, the internalization rate decreases, diminishing the difference in adhesion dynamics between the two cases.

**Fig. 8.**
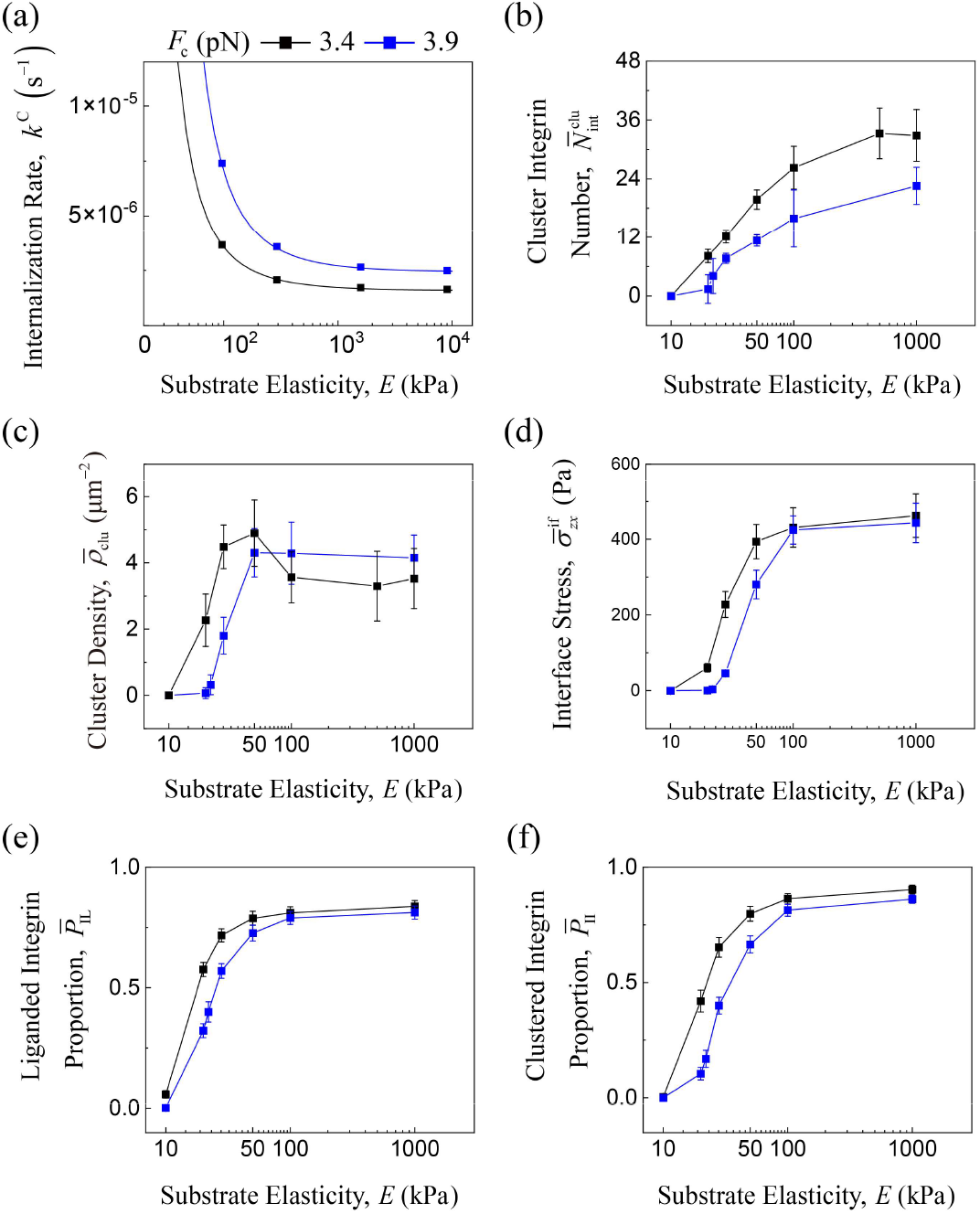
Effect of integrin internalization on the multiscale dynamics of cell–substrate adhesions: (a) The variation of integrin internalization rate with the Young’s modulus of the substrate under different internalization parameters; The mean value of (b) the number of molecules within adhesion clusters, (c) cluster density, (d) interface stress, and the proportion of (e) ligand-bound integrins and (f) clustered integrins at equilibrium state. In simulations, we take 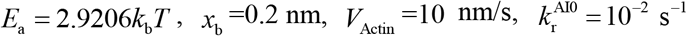 and *E* =3.9889 *k T*, considering constant rate II bonding processes.

Adhesion dynamics were analyzed on soft and medium-stiffness substrates (*E* =10–50 kPa) under three sets of internalization parameters. The relations between the internalization rate and the substrate stiffness is illustrated in Fig. 9, where the black line (*U*_C_ = −14.7 *k*_b_*T* and *F*_C_ =3.4 pN) and the red line (*U*_C_ = −15.5 *k*_b_*T* and *F*_C_ = 3.9 pN) intersect at approximately *E* = 68.5 kPa. To assess the formation of adhesion clusters, we consider the condition where 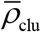 reaches 1 μm^−2^ and 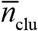 reaches 6.

**Fig. 9.**
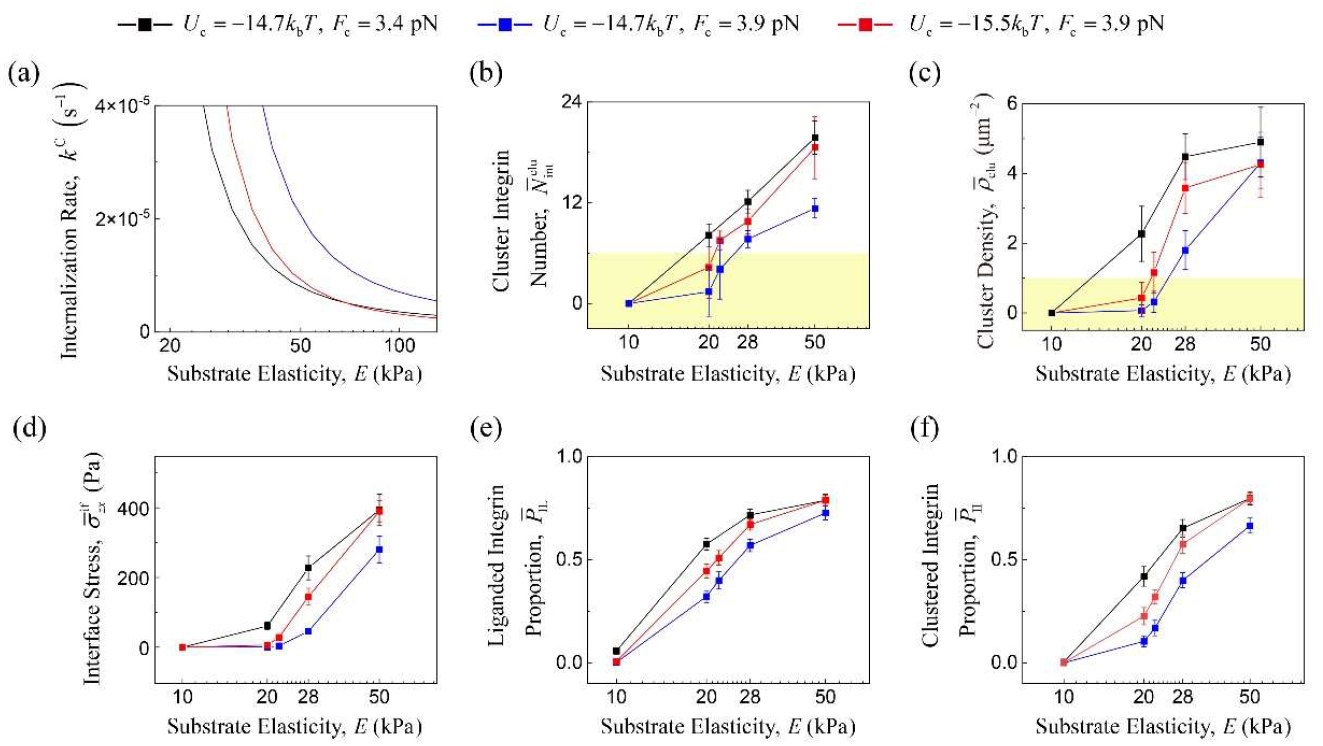
Effect of integrin internalization on the multiscale dynamics of cell–substrate adhesions when E = 20–50 kPa: (a) The variation of integrin internalization rate with substrate Young’s modulus under different internalization parameters; The mean value of (b) the number of molecules within adhesion clusters, (c) cluster density, interface stress, and the proportion of (e) ligand-bound integrins and (f) clustered integrins at equilibrium state. In simulations, we take 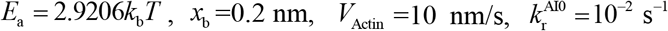 and *E*_II_ =3.9889 *k*_*b*_*T*, considering constant rate II bonding processes.

The intersection of the white and yellow regions in Fig. 9 (b, c) represents the threshold of the Young’s modulus 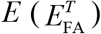. As shown in Fig. 9 (b, c), the case with the highest internalization rate (blue line) assumes 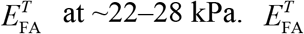 decreases to 20–22 kPa when the internalization rate decreases to the intermediate case (*U*_C_ = −15.5 *k*_b_*T* and *F*_C_ =3.9 pN), and further decreases to 10–20 kPa for the lowest internalization rate (black line). This finding suggests that a decrease in the internalization rate can lower the threshold of the Young’s modulus of the substrate required for the adhesion cluster formation.

### Multiscale mechanobiochemical adhesion dynamics

The interactions between the cell–substrate adhesion and the actin cytoskeleton are highly complex and dynamic, which play crucial roles in the regulation on adhesion dynamics and various cellular processes. However, fundamental questions, such as the spatiotemporal evolution of FAs under the influence of the flowing actin network, remain inadequately understood. To investigate the mechanical regulatory mechanisms of actin flow on FAs dynamics, this subsection will establish quantitative relationships between actin flow velocity and multiscale adhesion dynamics parameters through numerical simulations. To analyze the specific mechanical effects of actin flow, we consider a constant rate II binding process.

We simulate the spatiotemporal evolution of the cell–substrate adhesion under actin flow at speeds ranging from 0 to 20 nm/s on a substrate with *E*=100 kPa. As shown in Fig. 10(a), both of 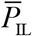 and 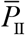 experience an approximately linear decline in the velocity range of 0 to 10 nm/s, followed by a gradual approach to equilibrium in the high-velocity regime (around 10 to 20 nm/s). These results indicate that the dissociation of IL and II bonds are promoted by actin flow, thereby inhibiting the ligand binding and integrin clustering. Despite the inhibition of adhesion by actin flow, the increase in the driving force from actin flow significantly enhances the transmission of traction forces at the cell–substrate interface [Fig. 10(c)]. The following equilibrium of force transmission reached in the high-velocity range can be attributed to the mechanosensitivity of IAC dynamics, where the failure and slippage of intermolecular force transmission pathways prevent the actin cytoskeletal forces acting on adhesions from further accumulating. Moreover, variations in actin flow velocity within the physiological range have little influence on the ratio of clustered molecules to liganded integrins 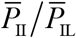 [Fig. 10(a)], whereas the substrate stiffness induces a pronounced effects on this ratio [Fig. 4(c)].

**Fig. 10.**
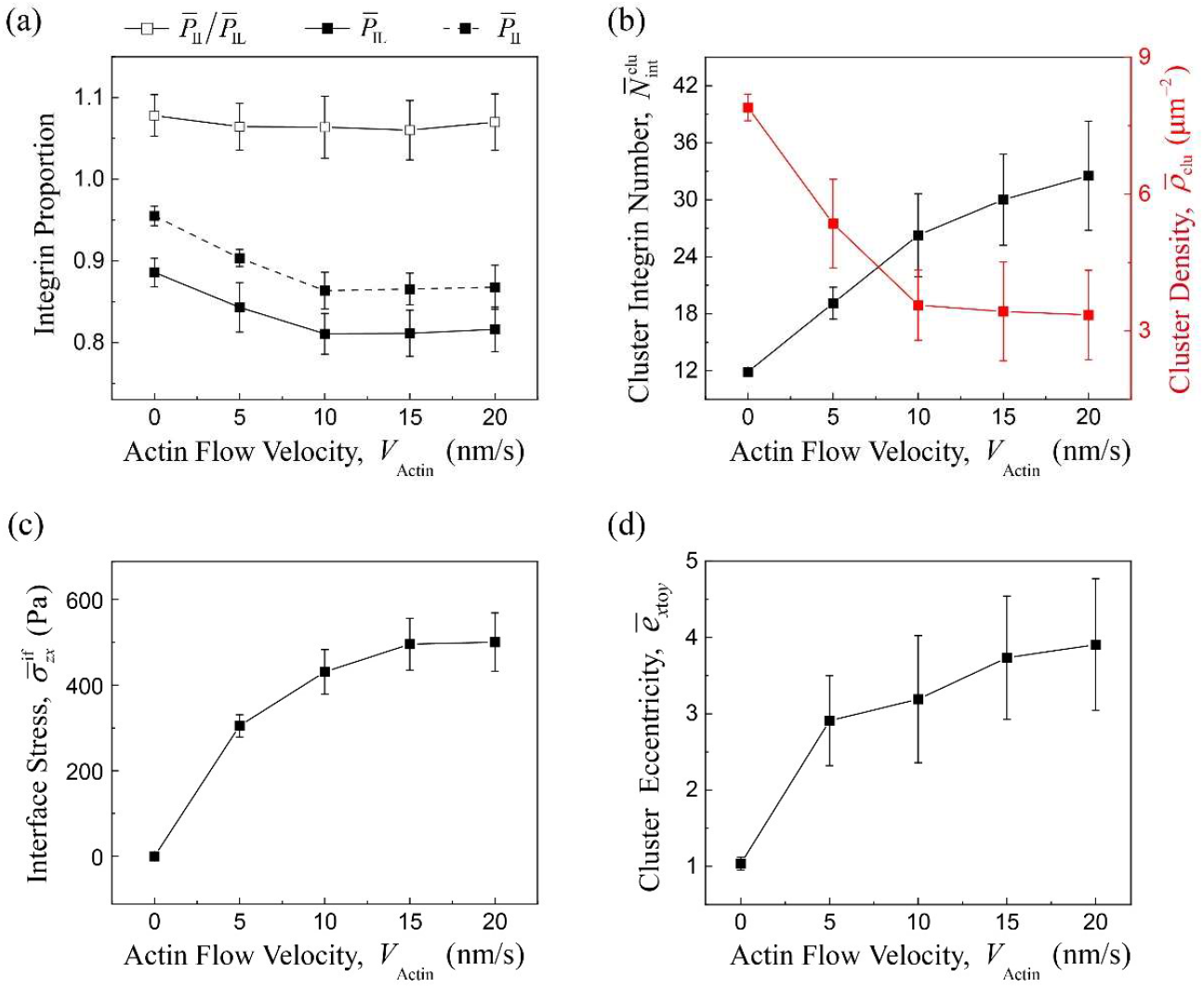
Effect of actin flow velocity on the multiscale dynamics of cell–substrate adhesions: the mean value of (a) the proportion of ligand-bound integrins and clustered integrins, (b) number of molecules within adhesion clusters and cluster density, (c) interface stress, and (d) cluster eccentricity at equilibrium state. In simulations, we take 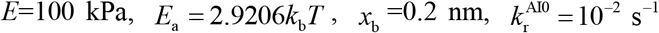, and *E*_II_ = 3.9889 *k*_b_*T*, considering constant rate II bonding processes.

The mechanical regulation of actin flow on FAs dynamics is shown in Fig. 10(b, d). As actin flow velocity increases, 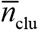 increases by nearly threefold, while 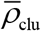 decreases by more about half [Fig. 10(b)]. This morphological transition from small and diffuse adhesions to large and sparse adhesions indicates that directed transport by actin flow can assist cell–substrate adhesions assembly into large and stable clusters. Specifically, the cytoskeletal force exerted on clustered integrins causes a part of adhesion clusters to disintegrate, allowing the released integrin molecules available for the further assembly of the remaining adhesion clusters. This is also consistent with the adhesion evolution process observed in previous experiments, where most nascent adhesions dissemble, and only about 20% of NAs eventually assemble into mature micron-sized FAs (55). This finding supports the promotion on FAs assembly by integrin transport, and shows that our model can reveal the physical effect of actin flow on the FA dynamics. Moreover, the directed transport by actin flow can also significantly enhance the anisotropy of adhesion clusters along the direction of shear flow [Fig. 10(d)] consistent with the results in our previous study without integrin internalization (35). These results reveal that the orientation of the promoted FA assembly is guided by the directed transport of the actin flow. In the simulation, the total number of integrin molecules is assumed to be a constant value. In vivo, however, signaling processes may induce variations in integrin concentration. These factors can be further explored in subsequent studies.

Then, the substrate stiffness, a key extra-cellular factor influencing the mechanical regulation of cell–substrate adhesion dynamics, is considered. To investigate its role, we conducted numerical simulations modulating substrate stiffness to examine its effect on multiscale adhesion dynamics mediated by actin flow velocity (Fig. 11). In this set of simulations, the characteristic length parameter *x*_b_ in Bell model takes 0.2 nm [Eqs. (16) and (17)]. As shown in Fig. 11(c), both actin flow velocity and substrate stiffness enhance the interfacial traction force, corresponds to a regime of appropriate mechanosensitivity that the force sensitivity of IAC bond rupture is at an appropriate level to ensure dependable force transmission. In the case of appropriate mechanosensitivity, the continuous gradient in number of molecules within adhesion clusters across the parameter space demonstrates a robust trend that actin flow and substrate stiffness cooperatively promote FA maturation (i.e., assembly) [Fig. 11 (b)]. According to the multiscale adhesion dynamics shown in Fig. 11, the coupled mechanical regulation mechanism of the actin flow and the substrate stiffness on adhesion morphology and force transmission behavior is summarized in the phase diagram shown in Fig. 12. As shown in Fig. 12(a), on a soft substrate with *E*= 10 kPa (region V), no obvious adhesion clusters can form due to the intense integrin internalization. When *E* increases to 20 kPa (region IV), sparse and small adhesion clusters can be observed. On stiffer substrates (*E*≥50 kPa), [i.e., regions I, II, and III in Fig. 12(a)], the integrin clustering is upregulated [Fig. 12(d)], then the actin flow can exert a more pronounced impact on adhesion morphology. As actin flow velocity increases, adhesion assembly on stiffer substrates is significantly promoted [Fig. 12(b)]. The adhesion morphology transitions from dense and small adhesions (region I) to medium-scale adhesions with medium density (region II) and then to low-density large adhesions (region III) [Fig. 12(a, b)]. Specifically, when *V*_Actin_ =0 and *E* =100–500 kPa, dense and small-scale adhesion form with 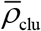 ranging around 7.9–8.2 μm^−2^ and 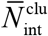 stable at ∼11.9, showing no significant change with increasing the substrate stiffness [Fig. 11(a, b)]. As *V*_Actin_ increases to 20 nm/s from 0, 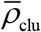 decreases by around two times, and 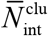 approximately increases threefold [Fig. 11(a, b)]. Under the actin flow with increasing velocity, the cell–substrate adhesions gradually evolve into stable FAs with large scale, a sparse distribution, and strong interface transmission capability. This morphological transition from small adhesion clusters to large FAs indicates that actin flow can promote adhesion assembly through directed transport on stiffer substrates. Interestingly, it is also noted that the FA assembly [Fig. 11(b)] is positively related to interface force transmission [Fig. 11(c)], rather than molecular scale adhesion activity [Fig. 11(d)], suggesting that effective interface force transmission relies on the formation of relatively stable higher-order adhesion structures.

**Fig. 11.**
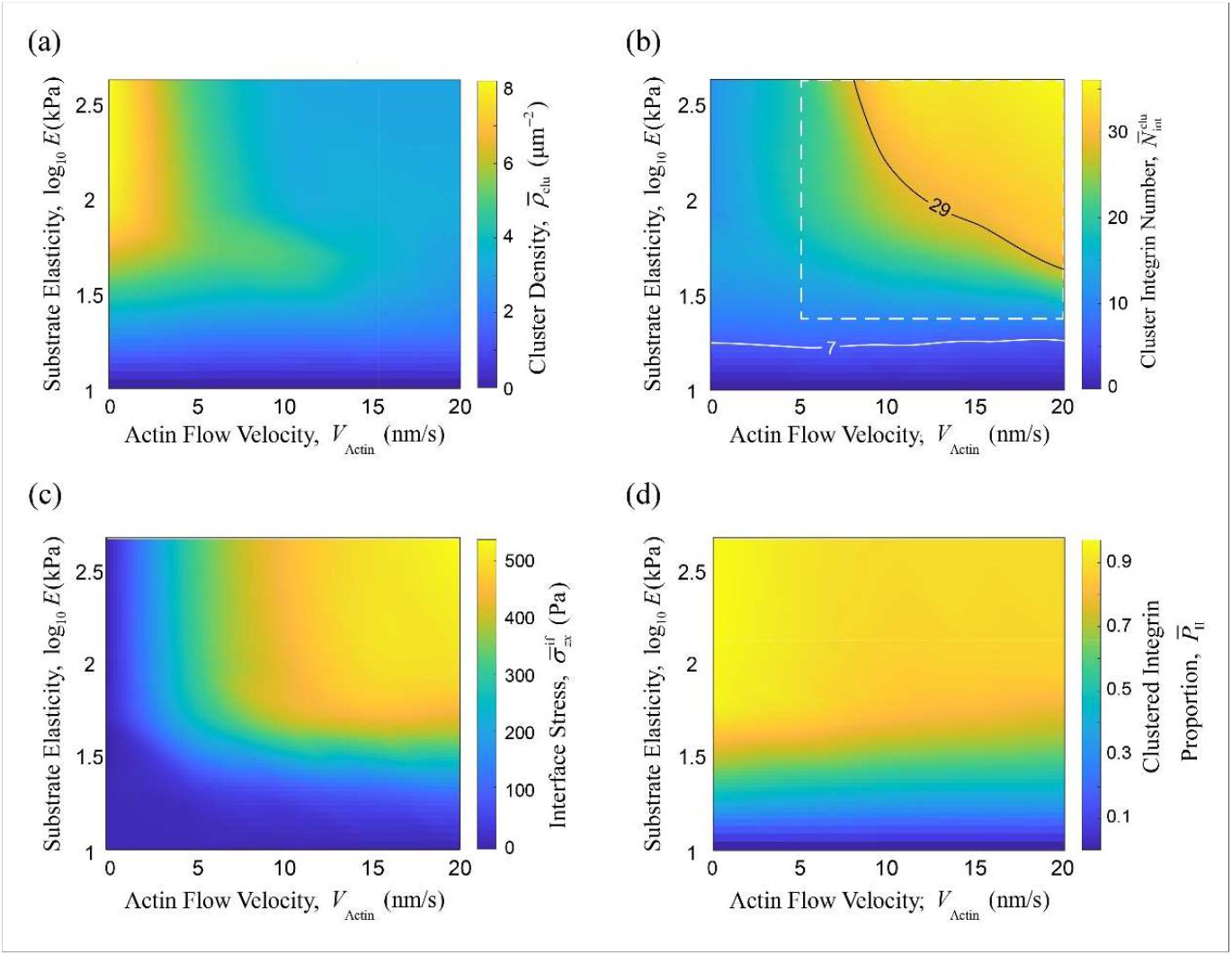
Mechanical regulation of the multiscale dynamics of cell–substrate adhesions by the actin flow velocity and the Young’s modulus of the substrate. The contour plots of the mean value of (a) cluster density, (b) number of molecules within adhesion clusters, (c) interface stress, (d) proportion of clustered integrins. In simulations, we take 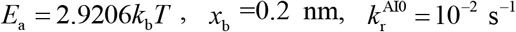, and *E*_II_ss = 3.9889 *k*_b_*T*, considering constant rate II bonding processes.

**Fig. 12.**
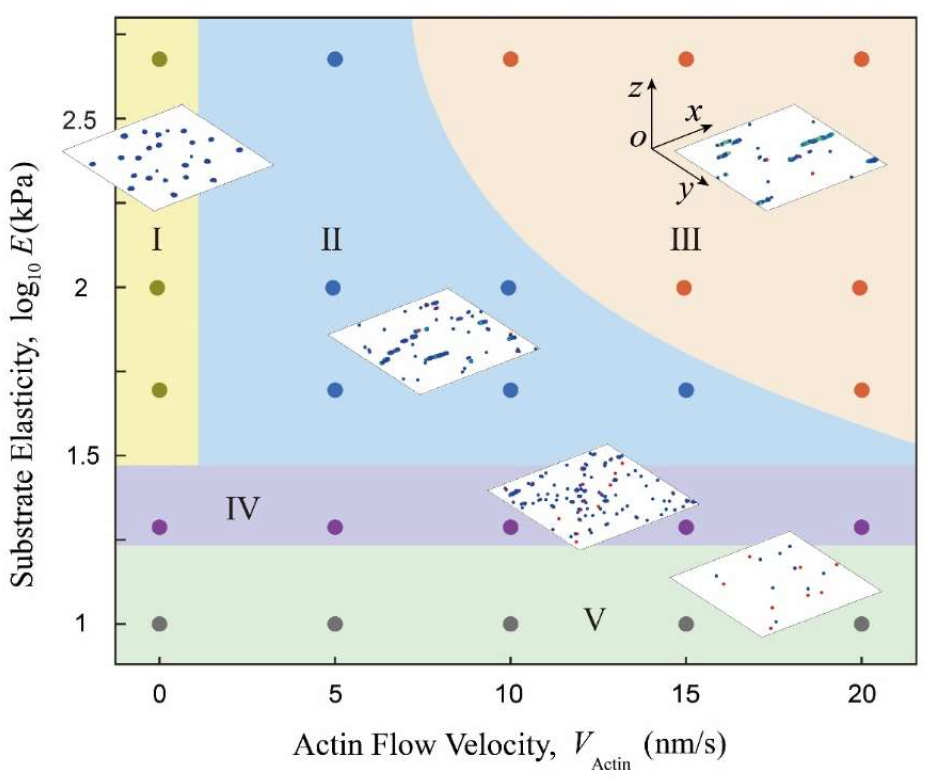
Mechanical regulation of the cell–substrate adhesion morphology by the actin flow velocity and the Young’s modulus of the substrate: the phase diagram of adhesion morphology, and representative adhesion morphologies in Region I–V. The parameters used here are the same as those in Fig. 11.

Although substrate stiffness shows no significant impact on the overall trend of FA remodeling by actin flow, it does modulate how readily FA maturation responds to changes in actin flow velocity. As illustrated by the 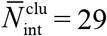 contour line in Fig. 11(b), stiffer substrates enable the same level of integrin assembly at lower actin flow velocities. For instance, to approach the assembly level of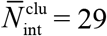, the required actin flow velocity decreases by more than half when increases from *E* 50 to 500 kPa. These results indicate that, on stiffer substrates, the promoting effect of actin flow transport on adhesion assembly becomes more pronounced. Then, we take a closer look into the failure mode of the IAC force transmission pathway within the white dashed box area in Fig. 11(b), as shown by Fig. 13(a). In the case of appropriate mechanosensitivity (*x*_b_ =0.2 nm), 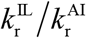 decreases with increasing substrate stiffness and actin flow velocity. The variation of 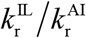 reveals that the dominant failure mode of the IAC force transmission pathway can transition from IL bond-mediated adhesion dissociation to AI interface slippage, facilitated by the increase of substrate stiffness and actin flow-mediated cytoskeletal forces [Fig. 13(b)].

**Fig. 13.**
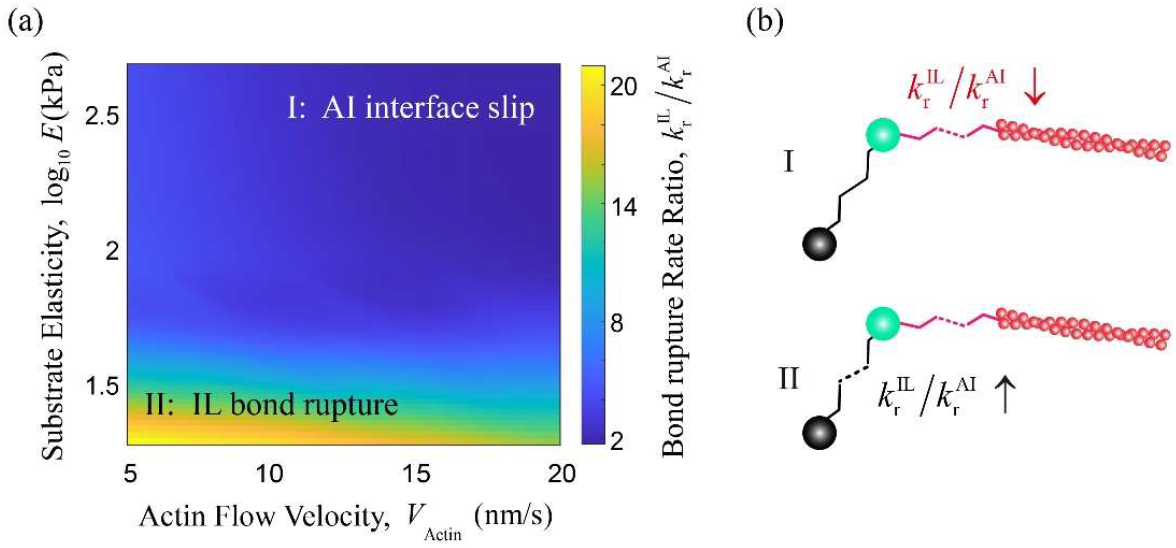
Failure mode of the IAC intermolecular force transmission pathway regulated by the actin flow velocity and the Young’s modulus of the substrate. (a) The contour plot of 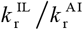. (b) Schematic of the failure mode of the IAC intermolecular force transmission pathway corresponding to Region I and Region II in (a). The parameters used here are the same as those in Fig. 11.

### Multiscale mechanobiochemical durotaixs mechanism

Cell–substrate adhesions are mechanosensitive subcellular structures embedded within a complex microenvironment. It connects externally to the ECM, and internally interacts with the cytoskeleton. Thus, its multiscale dynamics is regulated by the mechano-biochemical coupling mechanism driven by various extra- and intracellular microenvironmental factors. Among these factors, substrate elasticity and actin flow stand out as two key factors that have attracted significant scientific attention. In Subsection 4.3, numerical simulations considering the constant-rate integrin clustering process highlight the promoting role of actin flow in the directed physical transport of adhesion molecules. However, in vivo, the regulation of these microenvironmental factors on cell–substrate adhesions is closely related to downstream mechanotransduction steps, e.g., mechanosensitive conformational change of talin and vinculin mediated by their interaction with actin, leading to force-upregulated adhesion assembly [Eq. (18)]. Therefore, in this subsection, we analyze the multiscale adhesion dynamics regulated by substrate elasticity and actin flow transport, and examine the effect of downstream mechanotransduction on integrin clustering under actin flow.

To systematically investigate the coupling regulation by actin-driven integrin transport and substrate mechanics, we introduce two dimensionless quantities. First, the Péclet number *Pe* = *V*_Actin_ *L*_Scale_ *D* provides a dimensionless measure of actin-driven transport, quantifying the relative strength of actin-driven integrin transport versus stochastic diffusion. Here, *L*_Scale_ is a characteristic length of integrin cluster size, and is taken as 1 μm. Second, the relative substrate stiffness 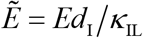 compares the elastic response of the substrate to that of individual integrin–ligand bonds, where *κ*_IL_ is Bond stiffness of IL. In numerical simulations, the actin flow velocity *V*_Actin_ and the Young’s modulus of the ECM *E* are set within the physiological ranges. To explore multiscale adhesion dynamics, the values of the characteristic length *x*_b_ are taken as 0.2 nm and 1.5 nm, representing two distinct levels of IAC bond rupture mechanosensitivity [Eqs. (16) and (17)], i.e., appropriate mechanosensitivity and high mechanosensitivity, respectively.

In the case of appropriate mechanosensitivity, the coupling regulation patterns of FA morphology and force transmission by relative substrate stiffness and actin-driven transport on are demonstrated through the phase diagram (Fig.14) and the contour plot of interface stress [Fig.15(a)]. As shown in Region I in Fig.14, almost no adhesion clusters form on soft substrates, so that the interface couldn’t bear any significant traction stress [Fig.15(a)]. On stiffer substrates 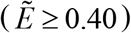 (i.e., Region II, III and IV in Fig.14), the actin-driven transport starts to show an increasing promotion on FA assembly and force transmission [Fig.15(a)] due to the weakened internalization and the consequently enhanced adhesion. As shown in Fig.14, sparse and small adhesion clusters form in the absence of actin-driven transport (Region II). As the actin-driven transport increases, the adhesion clusters initially transition to dense and medium-scale formations in Region III, and eventually evolve into large-scale clusters in Region IV. These trends align with the findings from Subsection 4.3, where similar effects on adhesion cluster formation are induced in cases of constant-rate integrin clustering. Comparing Fig. 11 with Fig.14, these results indicate that the incorporation of the force-regulated downstream mechanotransduction contributes to the actin-dependent adhesion cluster formation, in line with experimental cell observations (56–58).

In the case of high mechanosensitivity, the coupling regulation patterns of adhesion cluster morphology and force transmission by relative substrate stiffness and actin-driven transport are demonstrated by the phase diagram (Fig. 16) and the contour plot of interface stress [Fig.17(a)]. In this case, the enhanced sensitivity of IAC bond rupture engenders a biphasic regulation on adhesion assembly (about 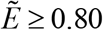) in response to both relative substrate stiffness and actin-driven transport (Fig. 16). Results in Region I show that adhesion clusters can hardly form on soft substrates, preventing effective force transmission [Fig. 16 and Fig.17(a)]. On stiffer substrates, small-scale adhesions in Region II can transition to large- or medium- scale adhesions in Region III or IV, respectively, as relative substrate stiffness or actin-driven transport increases. Subsequently, as either *Pe* or 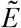 continues to increase, adhesion clusters in Region III or IV are further weakened and ultimately disassemble back into small- scale adhesions in Region II, contributing to the observed biphasic variations. Therefore, in this case, the assembly and nucleation of adhesion clusters peak at the interface with intermediate substrate stiffness and transport (Region III), leading to the formation of dense adhesion clusters with large scale. A biphasic interface stress variation with consistent trends of adhesion morphology is demonstrated by Fig.17(a). Moreover, it is noteworthy that, due to the negative regulation on adhesion assembly in the region with high actin-driven transport and relative substrate stiffness, further maturation of FAs fails when the mechanosensitivity of IAC bond rupture goes high.

**Fig. 14.**
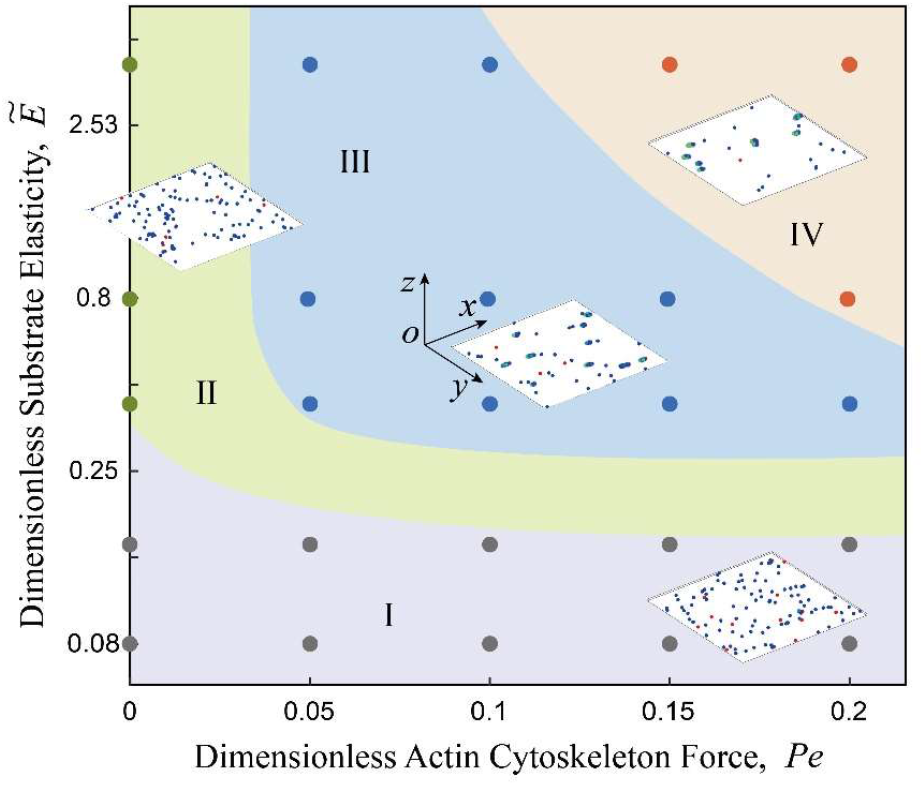
Regulation of cell–substrate adhesion morphology by actin-driven transport and substrate mechanics in the case of appropriate mechanosensitivity: I: no adhesion clusters form, II: medium-density, small-scale adhesions, III: high-density, medium-scale adhesions, and IV: medium-density, large-scale adhesions. In simulations, we take 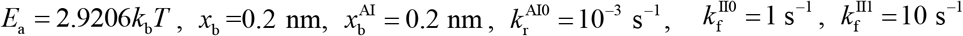, and *F*_b_ = 0.1 pN

**Fig. 15.**
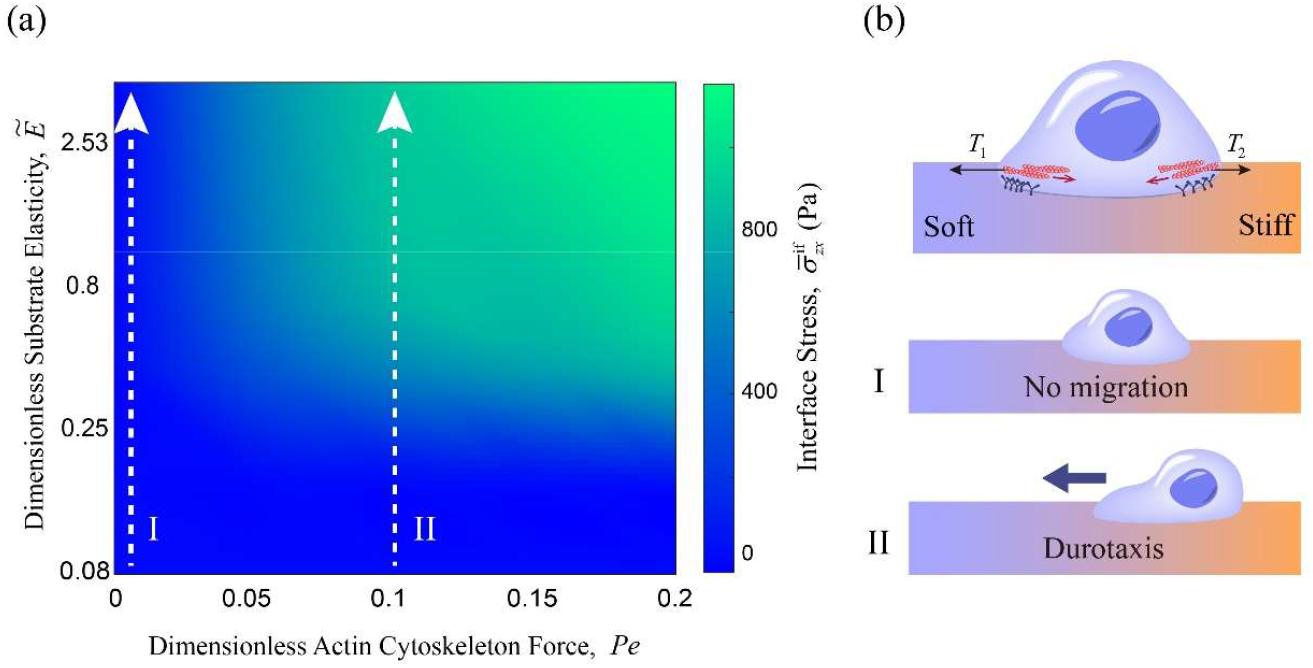
Regulation of cell–substrate interface stress and cell migration mode by actin-driven transport and substrate mechanics in the case of appropriate mechanosensitivity. (a) The contour plots of the mean interface stress at equilibrium state. (b) The schematic of cell migration state regulated by the gradient of substrate stiffness. Actin-driven transport magnitudes for cell migration modes I and II in (b) are illustrated by corresponding cases in (a), respectively. The parameters used here are the same as those in Fig.14.

**Fig. 16.**
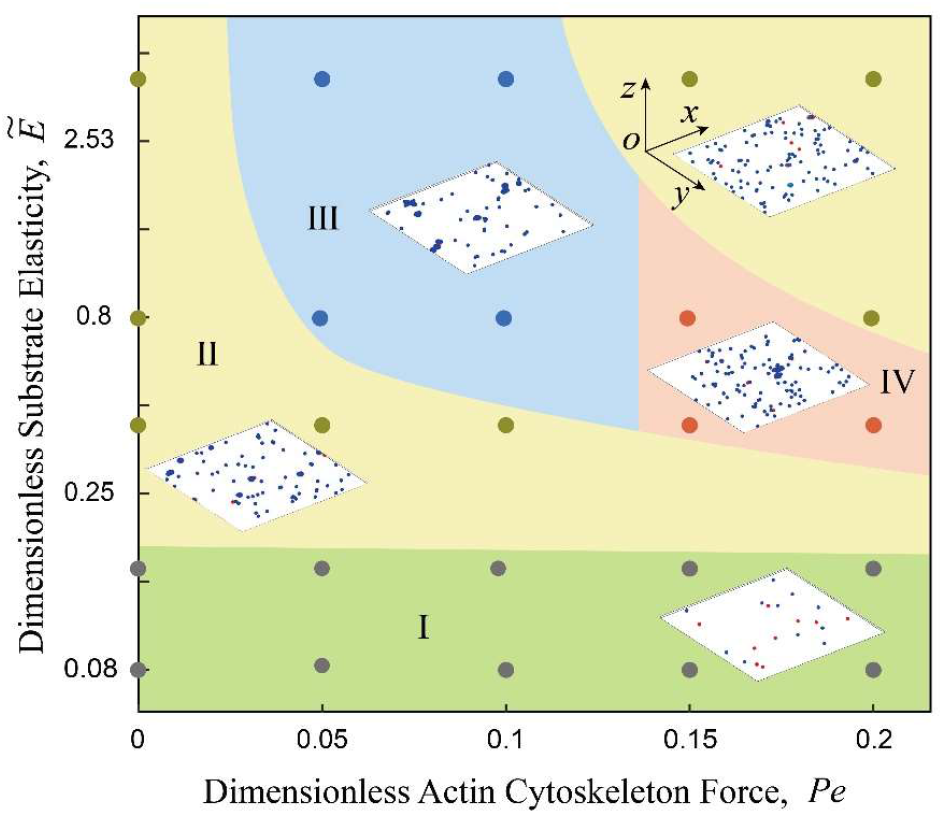
Regulation of cell–substrate adhesion morphology by actin-driven transport and substrate mechanics in the case of high mechanosensitivity: I: no adhesion clusters form, II: small-scale adhesions, III: high-density, large-scale adhesions, and IV: medium-scale adhesions. In simulations, we take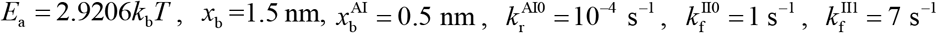, and *F*_b_ = 0.1 pN.

Furthermore, cell migration could be inferred from the traction dynamics at the cell–substrate interface. The interface stresses predicted by our model corresponds to those in the peripheral lamellipodia region on flat substrates. On flat substrates, adhesions at the cell–lamella interface are weak, and the cellular processes, e.g., spreading and migration, are primarily driven by the traction at the lamellipodia– substrate interface. During the cell spreading process, cell–substrate adhesions serve as anchoring points for the actin cytoskeletal traction system, which governs cell migration. Therefore, cell migration driven by the stiffness gradient of flat substrates could be revealed based on the predicted interfacial stresses.

As shown in Fig.15, in the case of appropriate mechanosensitivity, cell migration can hardly be driven by increasing relative substrate stiffness when the actin-driven transport is low [case I in Fig.15(b)]. However, as the actin-driven transport augments, cell migration could be driven by enhanced interface traction stress toward stiffer regions, i.e., durotaxis [case II in Fig.15(b)]. In the case of high mechanosensitivity [Fig.17(a)], weak interfacial stresses under low transport may not be sufficient to support cell migration cells either [case I in Fig.17(b)]. Subsequently, as the actin-driven transport increases, the effect of stiffer substrate on the cell migration mode turns to be durotactic [case II in Fig.17(a)]. In case III, a cell should exhibit durotactic migration when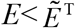, then transitions to negative durotaxis when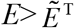, thus migrating toward the intermediate-stiffness region. This prediction suggests that the migrating cells may ultimately settle in regions with intermediate stiffness near the threshold. It is noted that the relative substrate stiffness threshold 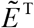, marking the transition from positive to negative correlation in the biphasic regulation of interface stress, tends to be negatively related to actin-driven transport when *Pe* is over ∼0.125 [case III in Fig.17(a)], suggesting that higher actin cytoskeletal forces could facilitate the reversal of cell migration direction by lowering the required substrate stiffness threshold.

**Fig. 17.**
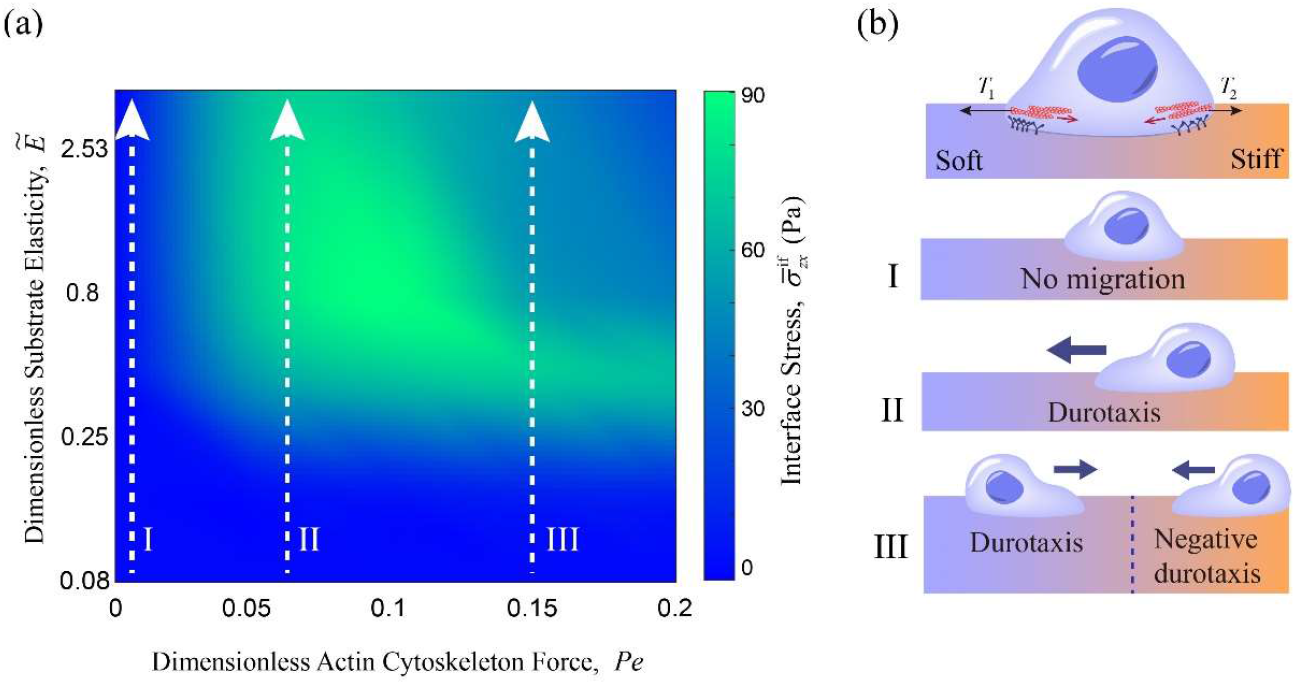
Regulation of cell–substrate interface stress and cell migration mode by actin-driven transport and substrate mechanics in the case of high mechanosensitivity: (a) The contour plots of the mean interface stress at equilibrium state. Actin-driven transport magnitudes for cell migration modes I, II and III in (b) are illustrated by corresponding cases in (a), respectively. The parameters used here are the same as those in Fig. 16.

## CONCLUSIONS

In summary, we have developed a multiscale dynamic model of cell–substrate adhesions to depict the mechanobiochemical coupling regulation process of cell adhesion. Specifically, our model incorporates integrin activation, clustering, and internalization to establish an integrin life cycle, considering mechanosensing activity mediated by endocytic carriers. It is worth noting that the caveolin-mediated integrin internalization pathway can be generalized to other mechanisms like clathrin-mediated pathways, as they both promote integrin vesicular trafficking by clustering and inducing membrane invaginations. Our model predicts that substrate rigidity inhibits integrin internalization, which in turn promotes integrin levels, FA assembly, and force transmission. With mechanical regulation of integrin internalization, the substrate rigidity sensing of the multiscale adhesion dynamics follows an S-shape rising trend. These results agree well with experimental measurements. We find an antagonistic effect between integrin internalization and integrin clustering governing the adhesion dynamics at molecule, FA and cell–substrate interface level. The cross-scale adhesion interactions play a crucial role in FA evolution, and can drive nonmonotonic regulation. Our results reveal a mechanobiochemical regulatory mechanism at the cell–substrate interface that influences FA morphology and cell migration modes, providing an explanation for durotaxis and negative durotaxis as observed in experiments.

Our model offers a theoretical framework for exploring the multiscale dynamics of cell–substrate adhesions, encompassing the comprehensive lifecycle of integrins. Therefore, it holds significant potential for elucidating the regulatory mechanisms at the cell–substrate interface that influence cross-scale processes involved in both physiological and pathological activities, including stem cell differentiation and cancer metastasis. For example, the model can examine how integrins orchestrate the adhesion, migration, and detachment of cancer cells by capturing cross-scale interactions, including integrin activities, FA dynamics, and cellular cytoskeleton reorganization. Furthermore, based on our model, the effects of the multiple interaction between molecular activities, e.g., integrin internalization and ligand binding, and gene expression regulation can be quantified, thus providing insights into the decision-making processes of stem cells regulated by mechanical and topological cues from the ECM.

## AUTHOR CONTRIBUTIONS

**H.L**.: Conceptualization, Methodology, Validation, Formal analysis, Visualization, Writing – original draft, Writing – review & editing. **W.F**.: Validation, Writing – review & editing. **X.C**.: Validation, Writing – review & editing. **B.L**.: Writing – review & editing. **X-Q.F**.: Supervision, Methodology, Formal analysis, Writing – original draft, Project administration, Writing – review & editing.

## CONTRIBUTIONS

Support from the National Natural Science Foundation of China (Grant Nos. 12032014, T2488101, and 12202248) are acknowledged.

